# Transcriptomic Analysis of *Plac1* Ablation Reveals Broad Alterations in Signaling Pathways Essential for Prenatal Development and Overlap with a Preeclampsia-Associated Signature

**DOI:** 10.64898/2026.04.30.721637

**Authors:** Suzanne Jackman, Xiaoyuan Kong, Yulan Piao, Alexei Sharov, Elin Lehrmann, Andrew Varshine, Ramaiah Nagaraja, David Schlessinger, Michael E. Fant

## Abstract

*Plac1* is an X-linked gene essential for placental and embryonic development. A knockout (KO) mouse model was used to define placental gene expression changes associated with Plac1 loss at E16.5 and E18.5 using gene expression microarray. Genes exhibiting at least a 1.5-fold change and FDR < 0.05 were considered significant. At E16.5, 717 genes were downregulated and 796 upregulated in KO placentas relative to wild type (WT), whereas at E18.5, 1121 genes were downregulated and 1151 upregulated. Subsets of highly and uniquely dysregulated genes were examined by gene-level curation alongside systems-level analyses, *Gene Ontology* (GO), *Kyoto Encyclopedia of Genes and Genomes* (KEGG), and *Ingenuity Pathway Analysis* (IPA), applied to the full differentially expressed gene (DEG) datasets. Downregulated genes were enriched for Rho GTPase-mediated and actin cytoskeleton-based processes, as well as membrane-associated signaling pathways with established roles in placental and embryonic development, vascular function, and branching morphogenesis. Overlap with pathways and molecular features associated with preeclampsia was also observed. In contrast, upregulated genes reflected, in part, immune activation and oxidative stress responses. These findings represent an important exploratory, hypothesis-generating analysis and provide a biologically coherent framework for understanding how Plac1 loss may be associated with placental dysfunction and pregnancy-related disease.

## 1. Introduction

*Plac1* (placenta-specific 1; human *PLAC1*) is an X-linked gene with high placental expression that plays essential role(s) in placental and embryonic development. *PLAC1* encodes a conserved, membrane-associated or extracellular protein expressed predominantly in trophoblast lineages, with localization near the apical region of differentiated syncytiotrophoblasts [1–5]. These features suggest a role at membrane-associated interfaces important for trophoblast function and maternal–fetal exchange.

An essential developmental role for Plac1 has been demonstrated in mutant mouse models. *Plac1*-null placentas exhibit placentomegaly, junctional-zone disorganization, trophoblast-lineage disruption, and mild fetal growth restriction [6,7]. Consistent with preferential paternal X chromosome inactivation in murine extraembryonic tissue [8,9], genetic analyses revealed that placentas derived from maternal (X^m-^X) heterozygotes (Hets) were phenotypically similar (but not identical) to knockout (KO) placentas whereas paternal (XX^p-^) Hets were phenotypically indistinguishable from wild type (WT). Surviving *Plac1*-null males and maternal heterozygotes also show increased susceptibility to develop postnatal hydrocephalus, suggesting that Plac1-dependent developmental programs also extend beyond the placenta [3]. Together, these findings support the view that Plac1 participates in biological processes required for normal placental development, fetal growth, and pregnancy maintenance.

PLAC1 has also attracted interest outside reproductive biology because it is reactivated in multiple cancers, where it has been associated with proliferation, migration, invasion, and immune modulation [10]. These observations do not establish equivalence between trophoblast biology and malignancy, but they support the broader concept that PLAC1 participates in conserved cellular programs involving membrane-associated signaling, tissue remodeling, and regulated cell–cell interactions.

Despite the established *Plac1*-null placental phenotype, the molecular pathways associated with the placental consequences of Plac1 loss remain incompletely defined. In particular, it is not known how Plac1 loss affects coordinated transcriptional programs involved in trophoblast organization, cytoskeletal regulation, vascular adaptation, immune modulation, and stress responses during late gestation.

In the present study, we re-examined placental transcriptomic data generated from *Plac1*-null and wild-type mouse placentas at E16.5 and E18.5, a developmental interval during which the mutant placental phenotype is well established [6]. The objective was not to define the complete mechanistic cascade downstream of Plac1, but to identify transcriptional patterns and pathway-level themes associated with Plac1 loss in the placenta. To this end, we integrated targeted genelevel curation with GO, KEGG, and IPA analyses.

Given the limited biological replication available from this legacy *Plac1* mutant colony, the findings are interpreted as exploratory and hypothesis-generating. Nonetheless, they support a model in which *Plac1* ablation is associated with broad alterations in signaling pathways essential for prenatal development, including those overlapping with molecular features associated with preeclampsia, and provide a biologically coherent framework for investigating how these perturbations may contribute to placental dysfunction and pregnancy-related disease within the Developmental Origins of Health and Disease framework.

## 2. Results

### 2.1 Developmental dynamics of placental growth in Plac1 mutants

This analysis was focused on the period of pregnancy when the growth trajectory of *Plac1*null placentas exhibit maximal divergence from WT placentas. Placentas associated with *Plac1*null embryos and maternal Hets (X^m-^X) exhibit placentomegaly and a disorganized junctional zone (JZ) [6]. Placental weights of X^m-^X Hets diverge from KO littermates between E16.5 and E18.5, likely because the paternal allele escapes complete inactivation. As we previously reported [6] and summarize in **Figure 1**, X^m-^X Het placental weight peaks at E16.5 and decreases slightly thereafter. By contrast, the weight of *Plac1*-null placentas continues to accelerate until E17.5 and then plateaus. These observations informed our decision to examine the developmental span bordered by E16.5 and E18.5.

**Figure 1.**
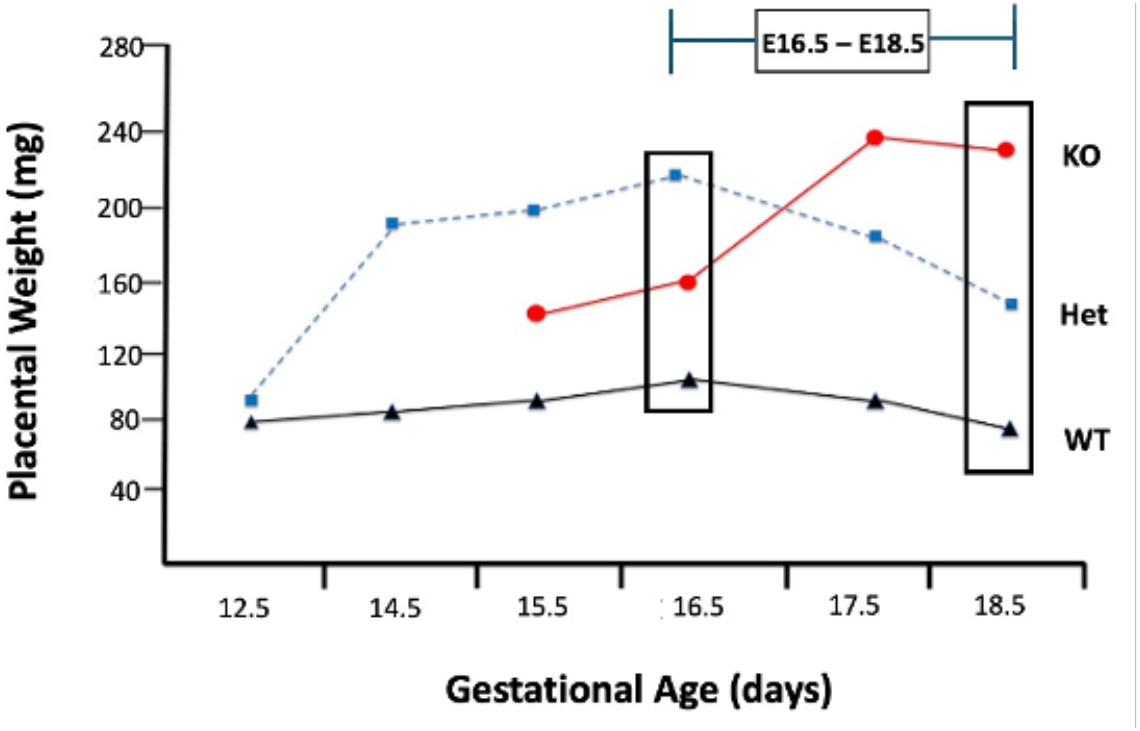
Growth trajectories of placentas from Plac1 mutants compared to WT placentas. Placental weights were determined throughout gestation representing WT, X^m-^X, and KO mice and presented in graphical form summarizing previously published data [6]. Each point represents the mean of age-specific samples for each genotype (n = 1-10 per data point). WT and KO data include both male and female placentas.

### 2.2 Gene Expression Microarray Analysis

The Agilent 4×44k gene chip was used to identify *Plac1-*dependent gene expression at E16.5 and E18.5. KO male placentas were compared to WT male placentas in duplicate samples. At E16.5, 717 known or putative genes were downregulated and 796 genes were upregulated at least 1.5-fold (FDR < 0.05) in KO placentas compared to WT. Similarly, at E18.5, 1121 genes were downregulated and 1151 genes were upregulated. (See supplemental materials, Tables S1S7, for processed data and complete gene lists).

Principal Component Analysis (PCA) (Supplemental Table S3) revealed that the first two components explained 67.03% of the variance (PC1: 41.7%, PC2: 25.33%). Visual inspection of PC1 and PC2 (**Figure 2**) indicates that PC1 primarily separated samples by gestational age (E16.5 versus E18.5, whereas PC2 distinguished genotypes (WT from KO).

**Figure 2.**
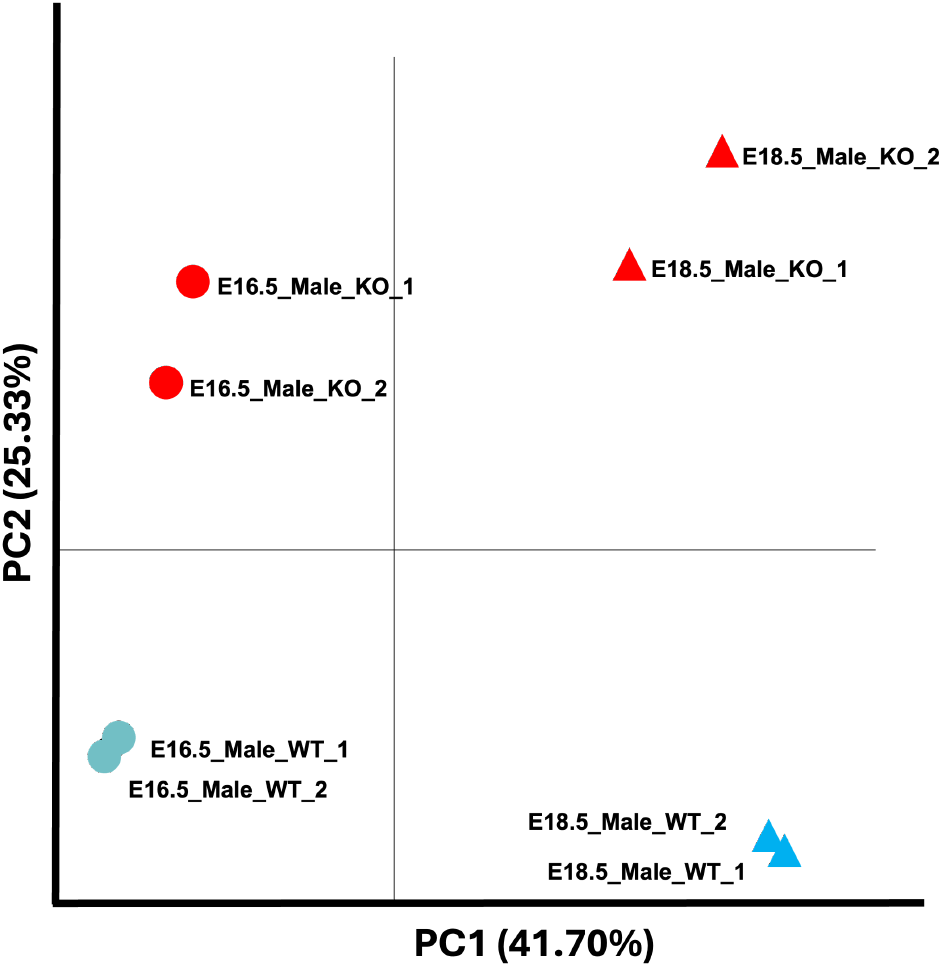
PCA of normalized expression data from E16.5 and E18.5 male placentas (WT and KO). PC1, and PC2 explain [41.70%] and [25.33%] of the variance, respectively. PC1 separates E16.5 vs E18.5, PC2 separates WT vs KO. Points are plotted by age, genotype, and replicate. **Color** = genotype (WT = blue, KO = red) **Shape** = age (E16.5 = circles, E18.5 = triangles)

KO versus WT scatter plots are shown for each age group (**Figure 3**). Most genes lie near the diagonal reference line, indicating broadly similar expression between KO and WT. Colored points denote differentially expressed genes (DEGs) meeting our threshold (≥1.5-fold change, FDR < 0.05). Red indicates upregulated genes in the *Plac1* KO and green indicates downregulated KO genes. Grey points indicate genes not significantly dysregulated. Axes show log₁₀ expression values.

**Figure 3.**
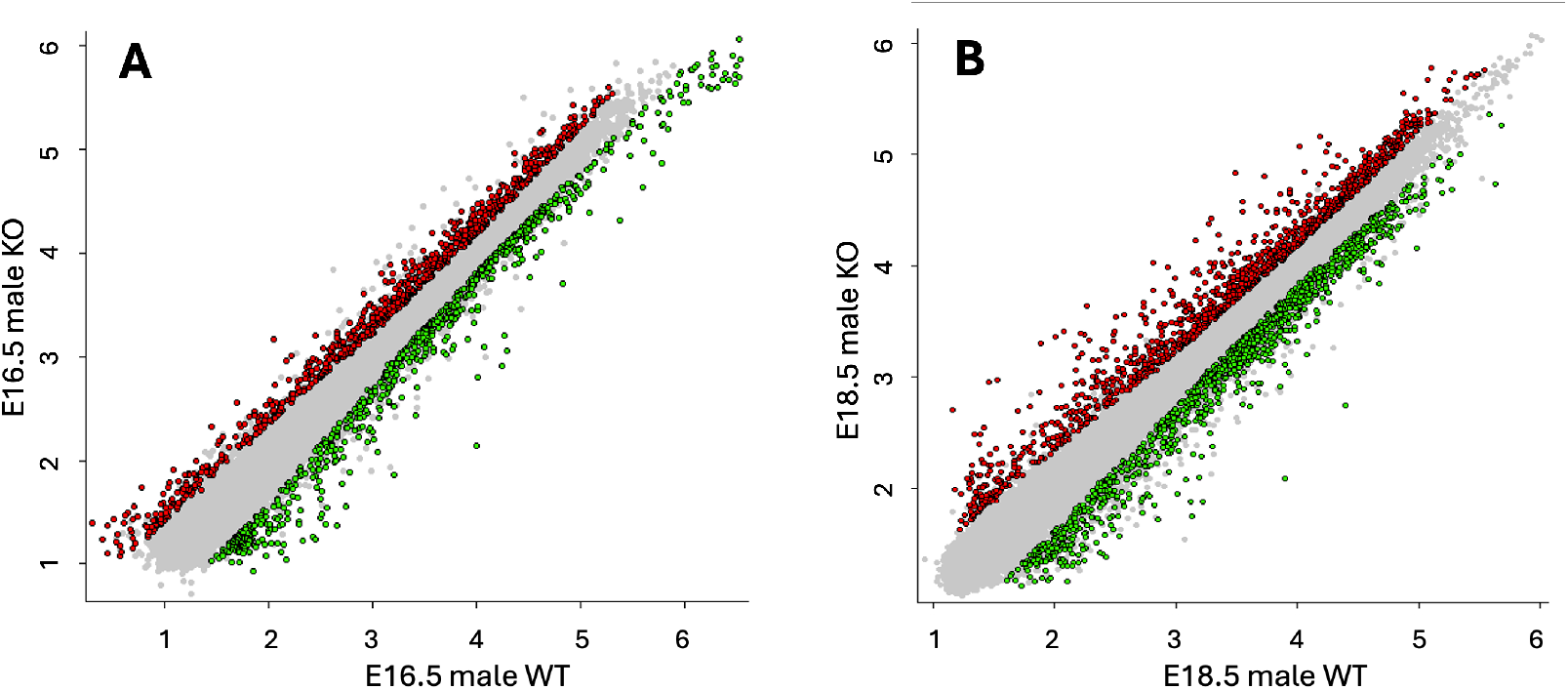
KO vs WT scatter plots by developmental stage. Pairwise scatter plots compare KO (yaxis) versus WT (x-axis) log10 expression within (**A**) E16.5 males (replicate means) and (**B**) E18.5 males (replicate means). Each point represents a gene/probe; the diagonal line indicates y = x (equal expression in KO and WT). Red points represent upregulated genes in the KO; green points represent downregulated genes in the KO; grey points represent no significant dysregulation, based on ≥1.5-fold change and FDR < 0.05

The heatmap of differentially expressed genes is shown in **Figure 4**. Expression values are visualized, with genes (rows) ordered by hierarchical clustering. qRT-PCR assays were performed at the time of the original microarray analysis for a limited panel of six genes at E18.5, with the number of assays constrained by colony attrition. These data are provided in Supplementary Figure S1 as qualitative directional observations only and not as independent validation of the microarray results.

**Figure 4.**
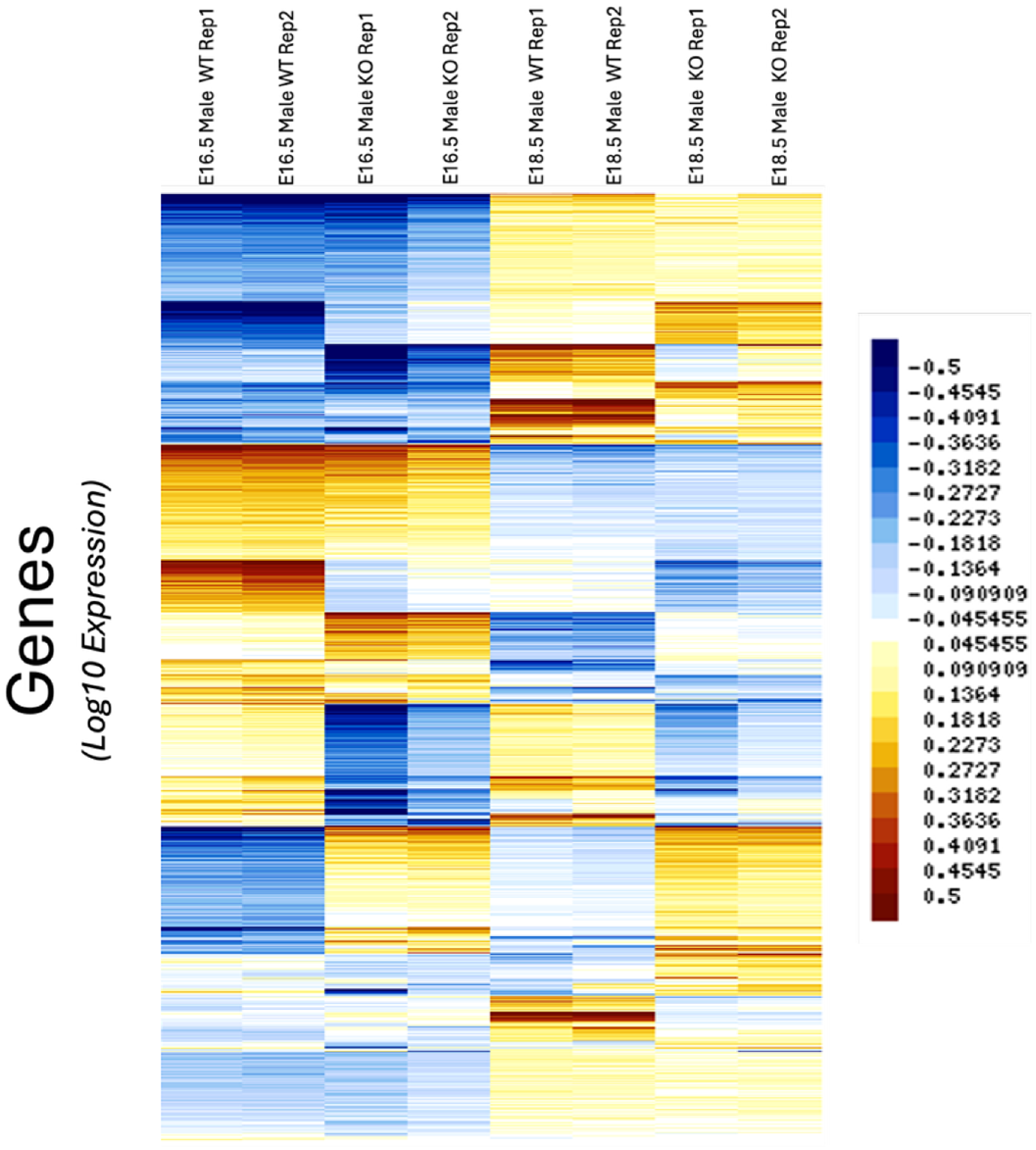
Heatmap of differentially expressed genes across samples. Heatmap displays relative gene expression patterns for differentially expressed genes identified at each developmental stage. For visualization, log_10_-transformed expression values were row-centered such that each gene’s mean expression across all samples equals zero. Color indicates expression relative to that gene’s mean level (blue, lower than average; yellow/brown, higher than average). Values were not variance-scaled. Genes (rows) are ordered by hierarchical clustering.

### 2.3 Gene-level Curation of Highly and Reciprocally Dysregulated Genes

We initially applied a targeted manual curation strategy to select subsets of differentially expressed genes (DEGs) to determine whether these transcripts could provide physiological context for the transcriptomic changes observed in the *Plac1*-null placenta. This approach was intended to be illustrative and hypothesis-generating rather than exhaustive, and to identify biologically coherent patterns that might otherwise be obscured when transcripts are considered only at the level of pathway enrichment.

Two complementary criteria were used for this purpose. First, we examined DEGs exhibiting the greatest magnitude of fold-change at each developmental stage (E16.5 and E18.5), irrespective of whether they were upregulated or downregulated (**Table 1**). This subset was considered because large-amplitude expression changes may reflect prominent biological responses within the *Plac1*-null placental environment. Second, we identified an additional subset of genes exhibiting reciprocal dysregulation across developmental stage, that is, genes downregulated at E16.5 and upregulated at E18.5 or, conversely, upregulated at E16.5 and downregulated at E18.5 (**Table 2**). These reciprocally regulated genes were considered because they may reflect evolving adaptive, compensatory, or decompensatory responses as placental dysfunction progresses during late gestation. (See Supplemental Tables S4–S7 for complete gene lists).

**Table 1.**
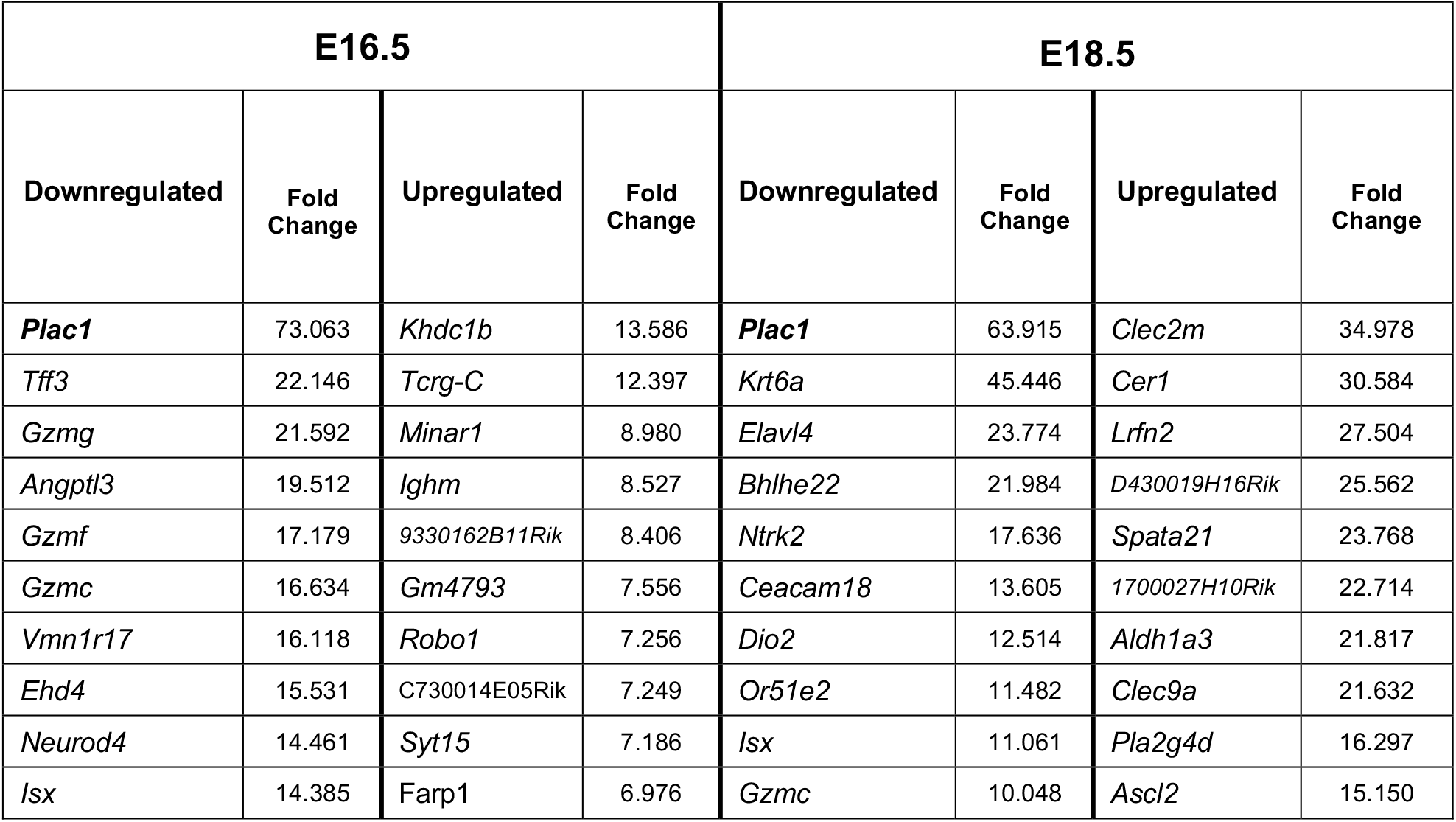
Genes Exhibiting the Greatest Dysregulation at E16.5 and E18.5. *(Fold-Change = Linear Scale; FDR < 0.05)*

**Table 2.** Genes Reciprocally Dysregulated at E16.5 and E18.5. *(Fold-Change = Linear Scale; FDR < 0.05)*

|  |  |  |  |  |  |
| --- | --- | --- | --- | --- | --- |
| **A. Dynamic Developmental Shift**<br>*(E16.5 Downregulated – to – E18.5 Upregulated)* |  |  | **B. Dynamic Developmental Shift**<br>*(E16.5 Upregulated – to – E18.5 Downregulated)* |  |  |
| **Genes** | **Fold Change**<br>**E16.5**<br>*(Downregulated)* | **Fold Change**<br>**E18.5**<br>*(Upregulated)* | **Genes** | **Fold Change**<br>**E16.5**<br>*(Upregulated)* | **Fold Change**<br>**E18.5**<br>*(Downregulated)* |
| *Asz1* | 2.412 | 2.537 | *Cnmd* | 2.715 | 3.766 |
| *Dnajb11* | 1.648 | 1.682 | *Crispld2* | 2.418 | 2.093 |
| *Dnajc3* | 1.834 | 1.582 | *Gpx3* | 2.748 | 2.012 |
| *Gm12618* | 1.891 | 1.604 | *Spon1* | 2.820 | 2.248 |
| *Gm773* | 3.337 | 2.159 | *Porcn* | 1.893 | 2.205 |
| *Noct* | 2.221 | 1.749 | *Tatdn2* | 2.430 | 2.165 |
| *Prl2c4* | 1.857 | 1.548 | *H6pd* | 2.003 | 2.591 |
| *Prl7a2* | 4.394 | 1.938 | *Siglecg* | 2.097 | 2.017 |
| *Prl8a8* | 1.710 | 1.995 | *Sphk1* | 2.080 | 2.344 |
| *Rab27a* | 1.797 | 2.143 | *1700064E03Rik* | 3.457 | 2.950 |
| *Tmsb10* | 2.877 | 1.703 |  |  |  |
| *Prom1* | 1.579 | 1.558 |  |  |  |

Considered collectively, the highest-magnitude DEGs at each time point (**Table 1**) clustered within several broad functional categories. These included genes associated with trophoblast structural integrity, membrane-associated signaling, cytoskeletal regulation, vascular development and endothelial interaction, immune modulation, and metabolic or stress-responsive processes.

Top-ranked downregulated genes included *Plac1* itself and genes associated with immune/trophoblast-interface signaling (*Gzmg, Gzmf, Gzmc*) [11–13], epithelial/cytoskeletal or adhesion-related programs (*Krt6a, Tff3, Ceacam18*) [14–16], membrane trafficking and receptor-mediated communication (*Ehd4, Ntrk2, Or51e2*) [17–21] vascular/endocrine-metabolic support (*Angptl3, Dio2, Isx*) [22–26], and developmental or neurodevelopmental annotations (*Elavl4*) [27]. Although *Neurod4*, *Vmn1r17*, and *Bhlhe22* were among the top-ranked downregulated transcripts, they lack convincing evidence for placental expression or function and were therefore excluded from consideration.

In contrast, upregulated genes were more often associated with immune/inflammatory signaling (*Tcrg-C, Ighm, Clec2m, Clec9a*) [28–30], metabolic or lipid-remodeling adaptation (*Aldh1a3, Pla2g4d*) [31, 32], and altered differentiation-, signaling-, or cellular remodeling-associated programs (*Minar1, Farp1, Ascl2, Cer1, Robo1, Syt15, Lrfn2*) [33–41]. Poorly characterized RIKEN or predicted transcripts were noted but not considered further. Interpreted at the level of functional clusters rather than as isolated gene-level effects, these patterns are broadly consistent with disruption of processes required for placental organization and maintenance and are generally concordant with the observed phenotype of *Plac1* mutant placentas.

The subset of reciprocally dysregulated genes (**Table 2**) provided an additional temporal perspective on the *Plac1*-null transcriptome. Genes shifting from reduced expression at E16.5 to increased expression at E18.5 included factors associated with ER/protein homeostasis (*Dnajb11*, *Dnajc3*) [42], vesicular trafficking or membrane-associated regulation (*Rab27a*, *Prom1*) [43, 44] and endocrine/placental lactogen-related support (*Prl2c4*, *Prl7a2*, *Prl8a8*) [45–47]. Conversely, genes elevated at E16.5 but reduced by E18.5 included factors associated with extracellular matrix or anti-angiogenic regulation (*Cnmd*, *Spon1*) [48–50], immune modulation (*Crispld2 [51]*, *Siglecg*) [52,53], oxidative/ER-redox stress handling (*Gpx3*, *H6pd*) [54–56], and lipid/signaling or developmental remodeling pathways (*Sphk1*, *Porcn*) [57–60].

These observations are consistent with altered temporal coordination of placental support programs in the absence of Plac1 and identify physiologically relevant transcriptional themes whose collective functional annotations align with the known *Plac1*-null placental phenotype. Importantly, this analysis does not assign causality or direct regulatory hierarchy to individual genes or pathways, nor does it establish mechanism. However, manual curation of these uniquely expressed gene subsets provides a complementary layer of analysis that preserves gene-level biological context when integrated with the pathway enrichment analyses.

#### 2.3.1 Functional Classes of DEGs

Examination of the DEGs also revealed a striking enrichment for membrane-associated receptors, solute transporters, and ion channels suggesting broad disruption of membrane-linked signaling and transport functions in the *Plac1*-null placenta. (**Table 3**). Notably, many of these genes are also annotated to brain development, consistent with shared placental and neurodevelopmental regulatory programs and with the CNS pathology observed in *Plac1* knockout mice [3].

**Table 3.**
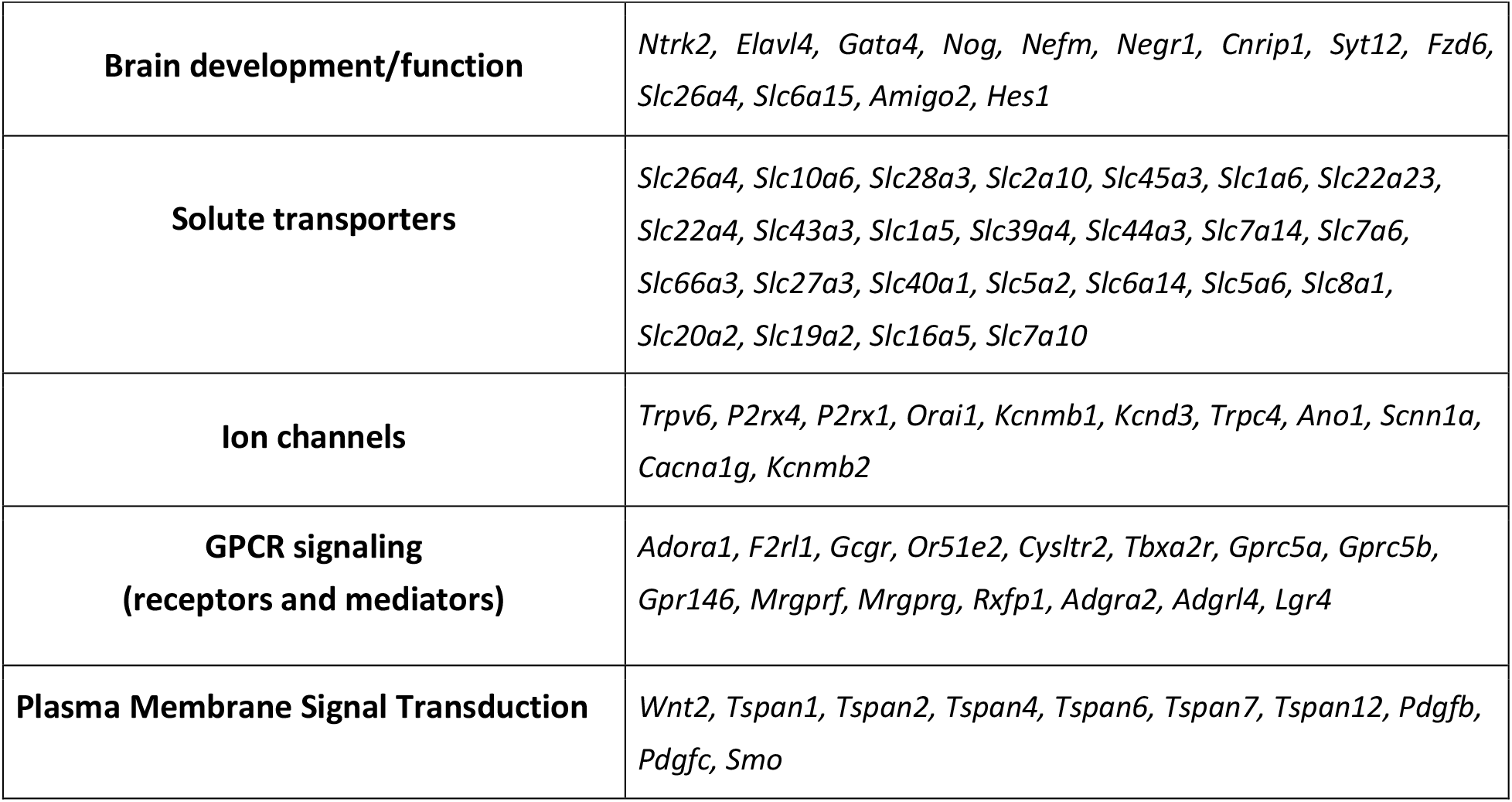
Functional Classes of Downregulated Genes (non-inclusive)

#### 2.3.2 Systems-level and Pathway Analyses: GO (Gene Ontology), KEGG (Kyoto Encyclopedia of Genes and Genomes), Ingenuity Pathway Analysis (IPA)

We next performed systems-level analyses to examine how transcriptomic changes associated with Plac1 loss converge on broader biological processes. GO, KEGG, and IPA analyses were used to identify functional themes and regulatory networks disrupted in the *Plac1*-null placenta, with emphasis placed on themes recurring across independent analytical frameworks.

##### GO Analysis

GO enrichment demonstrated predominant downregulation of cellular components (CC) associated with the membrane–cytoskeletal interface, including the apical, brush border plasma membrane, lysosomes, extracellular matrix, adherens and tight junctions, and actin-based structures at both E16.5 and E18.5 (**Figure 5 A, B**; see Supplemental Tables S8–S11 for complete GO term lists). These findings are consistent with the reported localization of PLAC1 to membranous compartments, particularly near the apical trophoblast surface. Additionally, enrichment of spindle-associated structures suggested possible effects on cytokinesis-related programs, while involvement of Weibel–Palade body-associated terms at E16.5 suggested altered endothelial-associated signaling. Given the role of Weibel–Palade bodies in the regulated release of von Willebrand factor, angiopoietin 2, and endothelin 1, genes associated with this compartment are particularly relevant to hemostasis, vascular tone, and angiogenesis, [61–63].

**Figure 5.**
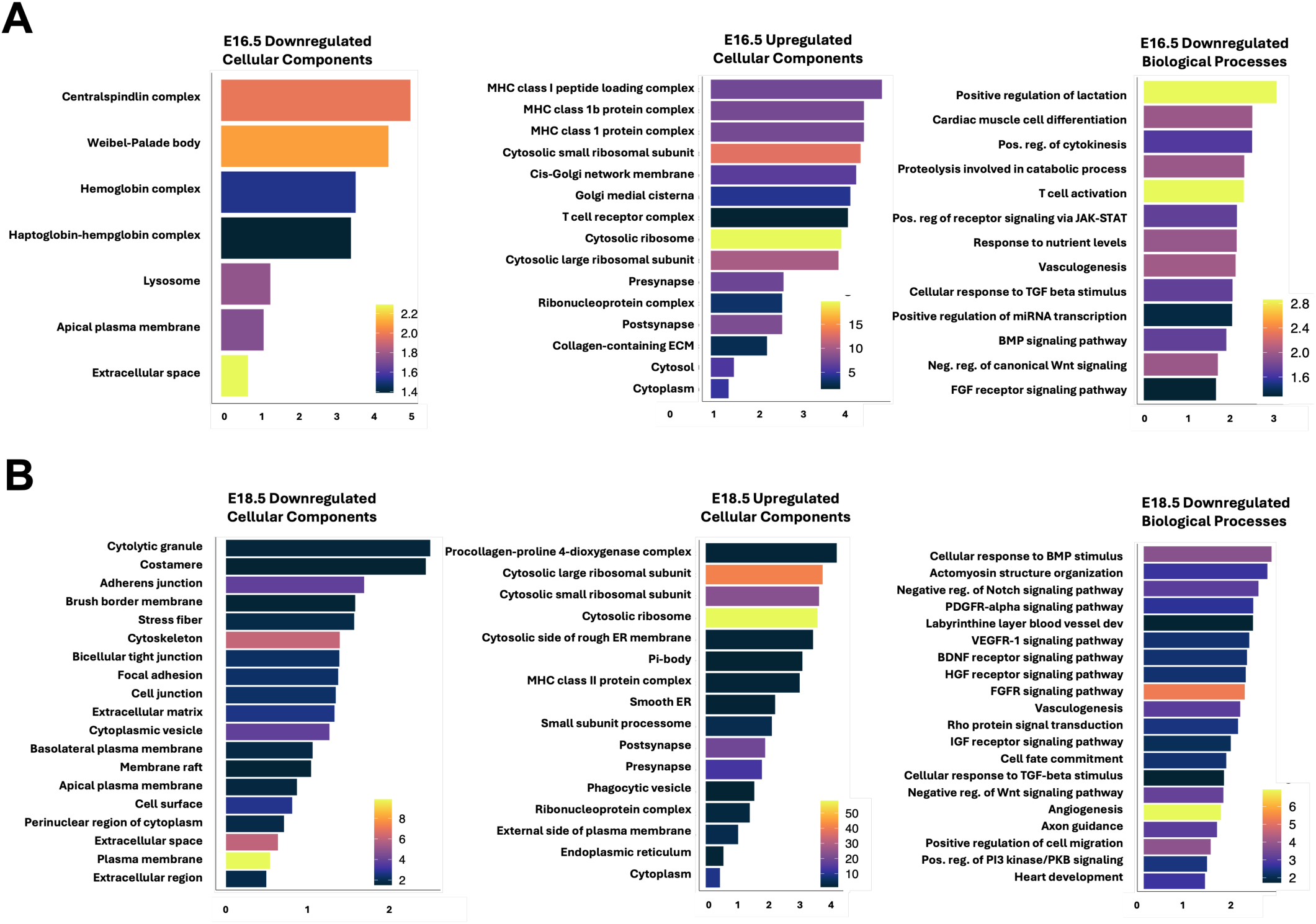
Developmental Stage-Specific Dysregulation of Cellular Components and Biological Processes in KO Placentas. Bars represent GO terms significantly enriched among the dysregulated genes. The X-axis shows log₂(Enrichment), indicating fold enrichment relative to WT controls. The Y-axis lists individual GO terms. Bar colors correspond to –log₁₀(FDR), with warmer colors (yellow/orange) indicating higher statistical significance and cooler colors (blue/purple/black) indicating lower significance. **Panel A:** E16.5, showing downregulated (left) and upregulated (middle) Cellular Components, and downregulated Biological Processes (right). **Panel B:** E18.5, showing downregulated (left) and upregulated (middle) Cellular Components, and downregulated Biological Processes (right). * Note – For visualization purposes, the number of BPs and CCs were selectively limited to a maximum of 20 by reducing implied functional redundancy.

In contrast, upregulated CC terms reflected stress-adaptive and compensatory responses, most notably enrichment of translational machinery, ribonucleoprotein complexes, and ER/Golgi-associated compartments, particularly at E18.5. These changes are consistent with increased translational load, secretory activity, and proteostasis demand under nutrient and oxidative stress. Immune-related components also showed significant enrichment, represented by the MHC class I complex at E16.5 followed by MHC class II complex and phagocytic vesicles at E18.5. Additional enrichment of extracellular matrix and synapse-annotated compartments, likely reflecting vesicle trafficking and cytoskeletal remodeling, further supported broad alteration of membrane dynamics and cell–cell communication.

Downregulated GO biological process (BP) analysis revealed enrichment of growth factor and developmental signaling pathways, including BMP, FGF, Wnt, TGFβ, and Jak–STAT, all of which have established roles in placental development [64–66]. In addition, dysregulated genes in the placental dataset were enriched for GO annotations related to embryonic organogenesis, particularly cardiovascular pathways. These annotations likely reflect shared developmental signaling programs used across embryonic tissues, including placenta, rather than direct evidence of organ-specific developmental pathology within placental tissue. GO BP enrichment for upregulated genes at both E16.5 and E18.5 was represented by generalized immune, stress, and metabolic response terms that overlapped extensively with KEGG and IPA pathway annotations described below and is presented in full in the Supplementary Data (Tables S9 and S11).

##### KEGG Analysis

KEGG analysis at E18.5 reinforced these findings, identifying enrichment among downregulated genes for pathways related to vascular smooth muscle contraction, calcium and cGMP–PKG signaling, tight junction integrity, TGFβ signaling, and Hippo pathway components (**Figure 6**; see Supplementary Tables S12–S15 for complete KEGG term lists). These pathways are broadly associated with vascular regulation, cytoskeletal organization, trophoblast proliferation, and structural stability, and are therefore consistent with altered placental remodeling and maternal–fetal interface function. No significant downregulated KEGG terms were identified at E16.5.

**Figure 6.**
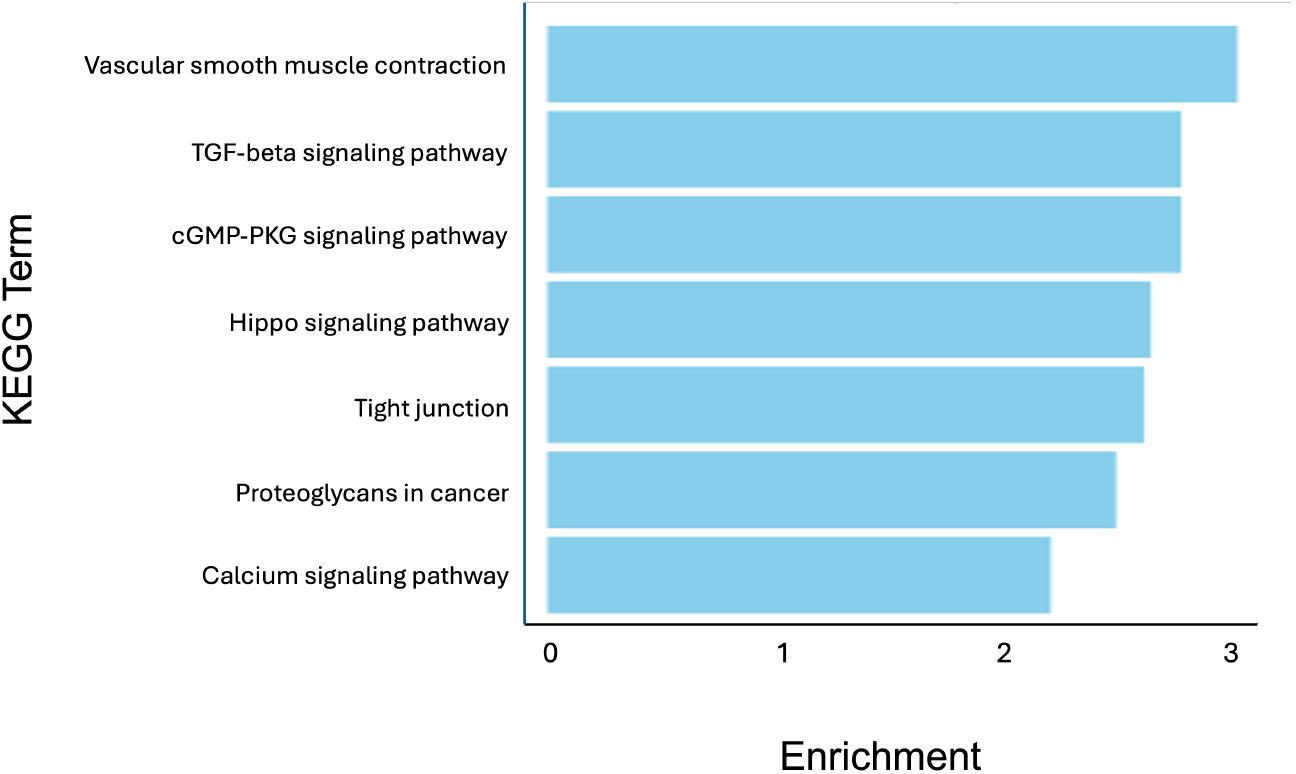
Fold enrichment of KEGG terms represented by downregulated genes at E18.5. Bars represent KEGG pathways significantly enriched among downregulated genes at E18.5. The X-axis shows fold enrichment, and the Y-axis lists individual KEGG terms (FDR < 0.05).

Conversely, KEGG terms enriched among upregulated genes were dominated by cellular stress, metabolic adaptation, and immune activation, including Ribosome, Protein Processing in the Endoplasmic Reticulum, Glycolysis/Gluconeogenesis, Phagosome, and Antigen Processing and Presentation (**Table 4**; Supplementary Tables S13 and S15). The viral pathway annotation, *Coronavirus Disease – COVID 19*, likely reflects heightened innate and adaptive immune signaling rather than pathogen-specific responses.

**Table 4.** Curated KEGG Terms Associated With Upregulated Genes.

| KEGG Term | E16.5 Fold Enrichment | E16.5 FDR | E18.5 Fold Enrichment | E18.5 FDR |
| --- | --- | --- | --- | --- |
| Ribosome | 6.63 | 3.82E-17 | 9.28 | 8.62E-45 |
| Coronavirus Disease-COVID 19 | 5.07 | 3.60E-13 | 6.81 | 1.44E-35 |
| Glycolysis/Gluconeogenesis | 5.82 | 2.77E-04 | --- | --- |
| Antigen processing/presentation | 5.65 | 3.28E-05 | --- | --- |
| Allograft rejection | 6.74 | 3.28E-05 | --- | --- |
| Graft versus host disease | 6.74 | 3.28E-05 | --- | --- |
| Biosynthesis of amino acids | 4.11 | 1.55E-02 | --- | --- |
| Pentose phosphate pathway | 5.65 | 2.50E-02 | --- | --- |
| Amino and nucleotide sugar metabolism | 5.75 | 1.55E-02 | --- | --- |
| Phagosome | 2.27 | 2.98E-02 | --- | --- |
| Cell adhesion molecules | 3.09 | 8.41E-03 | --- | --- |
| HIF-1 signaling pathway | 3.40 | 2.13E-02 | --- | --- |
| VEGF signaling pathway | 4.42 | 2.50E-02 | --- | --- |

##### Ingenuity Pathway Analysis (IPA)

Broadening our systems-level interpretation of the *Plac1*-null transcriptome, we next applied IPA analysis. Unlike gene-level fold-change comparisons, IPA integrates both the directionality of gene expression changes and curated regulatory relationships among pathway components to predict pathway activity states. Canonical pathways were therefore evaluated using the IPA activation Z-score, which estimates whether the collective expression pattern of pathway-associated genes is more consistent with activation or inhibition relative to random expectation. This approach enables functional inference at the pathway level and complements the enrichment-based GO and KEGG analyses described above.

IPA “Comparison Analysis” identified Rho GTPase signaling as the most significantly downregulated pathway across both developmental stages (**Figure 7A**; see Supplementary Tables S16–S19 for complete pathway lists), along with myocardin signaling, elastic fiber formation, membrane repair, cardiomyocyte differentiation via BMP receptors, and embryonic stem cell pluripotency pathways [67–69]. Upregulated IPA pathways were highlighted by predicted activation of translational control, ribosomal quality control, nonsense-mediated decay, EIF2/GCN2-mediated integrated stress response and antioxidant metabolism, consistent with chronic nutrient and/or oxidative stress [70–72] (**Figure 7B**; see Supplemental Table S20 and Figure S2). Suppression of GAIT and coronavirus pathogenesis pathway annotations may reflect altered inflammatory and antiviral-response regulatory programs, including impaired translational restraint of cytokine-associated signaling [73,74]. Collectively, GO, KEGG, and IPA analyses converge on a model in which loss of *Plac1* disrupts membrane-proximal, actin-cytoskeletal, and vascular regulatory networks, accompanied by progressive placental stress, immune dysregulation, and functional decline.

**Figure 7.**
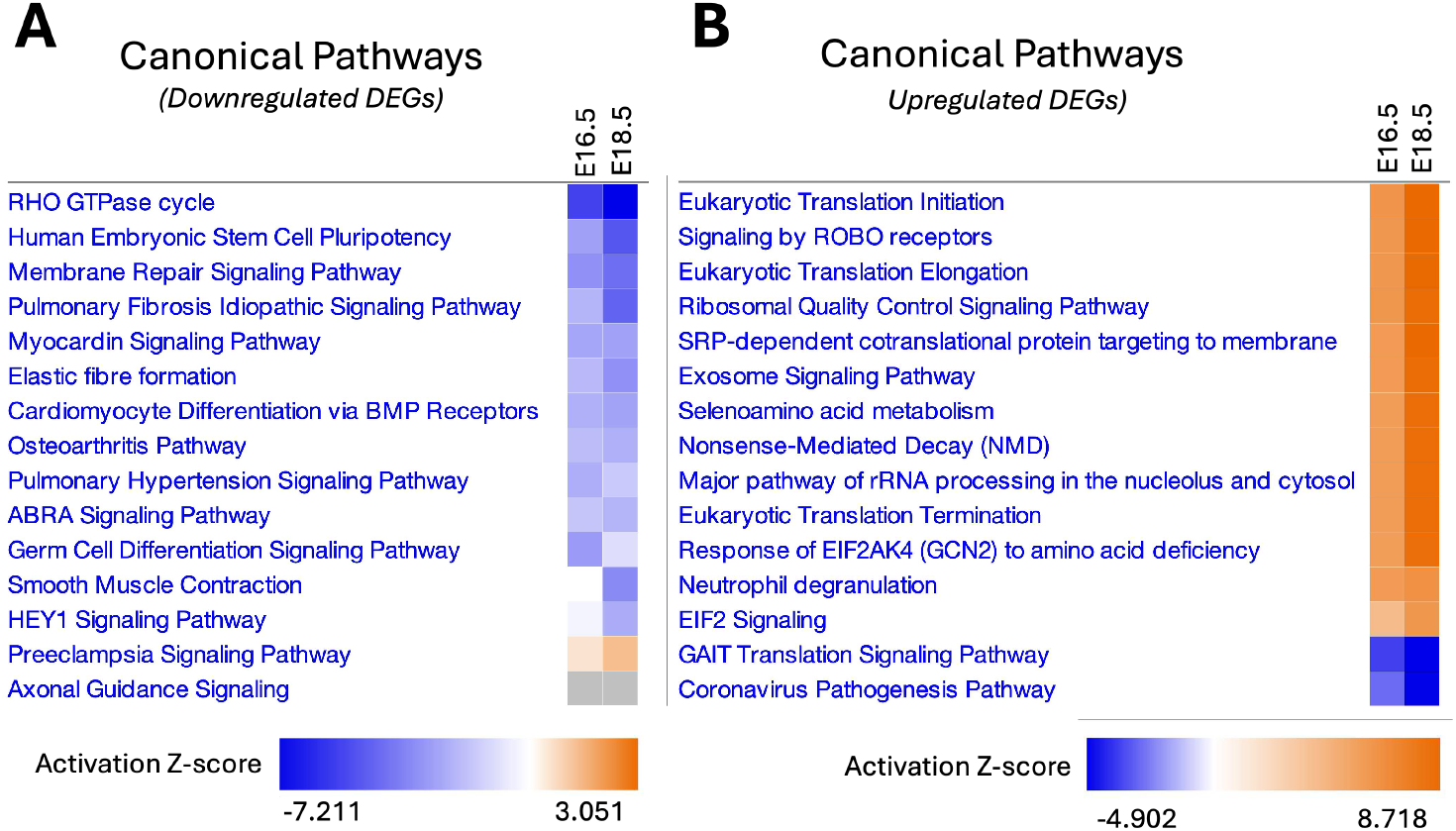
Heatmaps Depicting Developmental Stage-Specific Dysregulation of IPA Canonical Pathways in KO Placentas. **Panel A.** Canonical Pathways associated with Downregulated genes at E16.5 and E18.5. **Panel B.** Canonical Pathways associated with Upregulated genes at E16.5 and E 18.5. Color intensity reflects activation Z-scores: blue indicates negative Z-scores (predicted inhibition), and orange indicates positive Z-scores (predicted activation). Pathways filtered for absolute Z-scores > 2 and Benjamini–Hochberg adjusted p-values < 0.05 at one or more developmental stages and ranked by absolute Z-score

#### 2.3.3 Downregulation of Developmental Signaling Annotated to Brain and Cardiovascular Systems

Several IPA canonical pathways associated with downregulated DEGs were annotated to developmental processes involving cardiovascular and neural systems (Supplemental Tables S21 and S22). Cardiovascular-associated pathways were prominently represented during this timeframe, becoming more significant as gestation approached E18.5. Pathways meeting our statistical thresholds included Rho GTPase signaling, myocardin signaling, cardiomyocyte differentiation via BMP receptors, Hey1 signaling, and ABRA signaling (**Table 5**) [67–69]. Pathways annotated to brain development were also enriched, including Rho GTPase and axonal guidance signaling. Rho GTPase pathways have established roles in CNS development and function, and axonal guidance pathways contribute to branching morphogenesis [75–77], a process relevant to placental development and other embryonic tissues. This was notable given the known CNS phenotype we previously reported for this mouse model [3]. These annotations are reported as pathway-level outputs of the placental transcriptomic analysis and are interpreted as shared developmental signaling programs rather than direct evidence of organ-specific pathology.

**Table 5.**
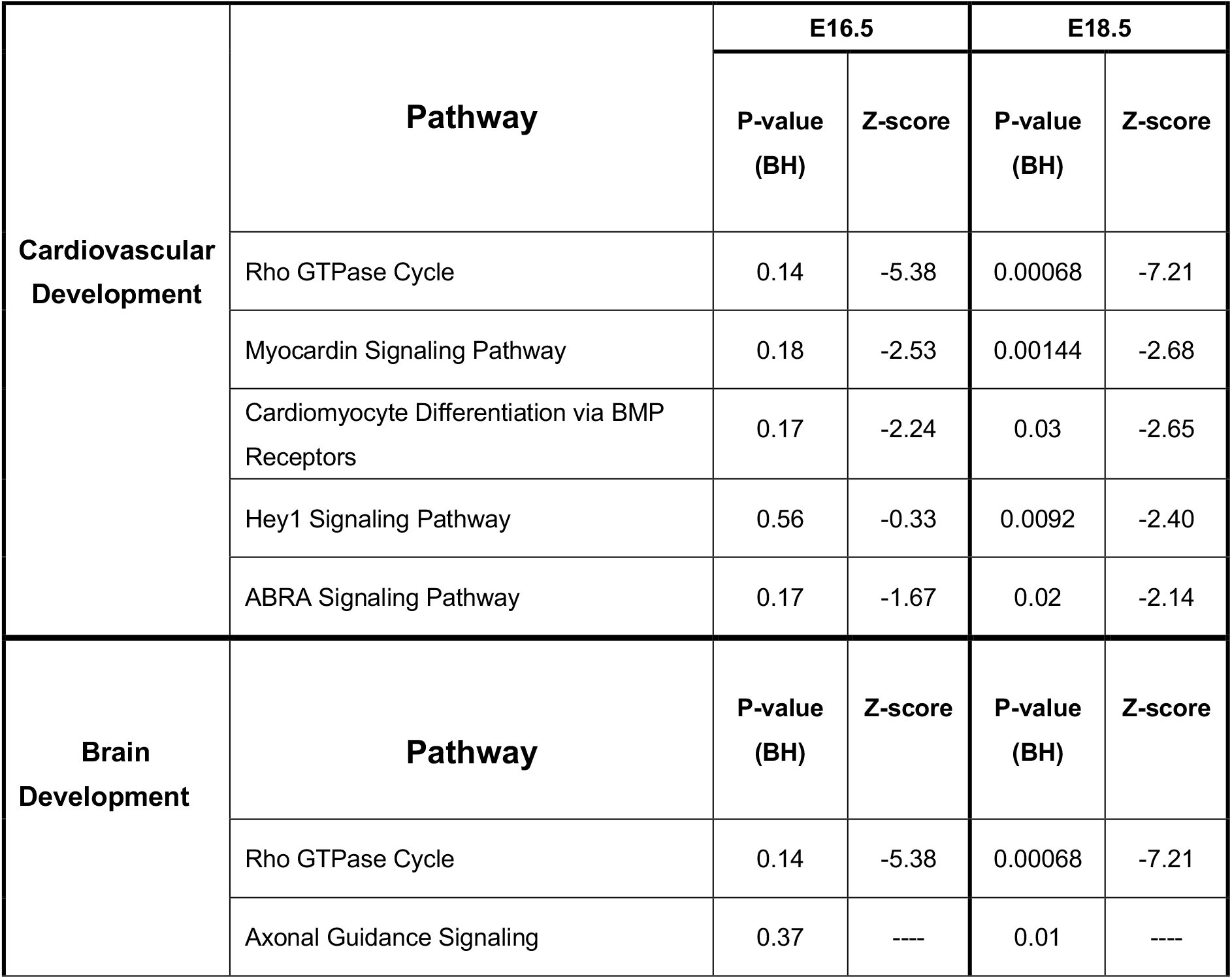
Canonical Pathways Associated with Cardiovascular and Brain Development. *(Filtered for B-H-adjusted p-value < 0.05 at one or more gestational ages)*

#### 2.3.4 Transcriptomic Overlap with Preeclampsia-Related Pathways and Molecular Features

The *Plac1*-null transcriptome showed overlap with pathways and molecular features reported for preeclampsia (PE) and related placental stress states. DEGs contributing to the PE-associated canonical pathway were identified at E16.5 and showed increased pathway-level concordance at E18.5 (**Figure 8**). At E18.5, this pathway reached statistical significance (FDR = 0.02) and showed a positive activation Z-score shift from 0.58 to 1.34. Because the Z-score did not reach the conventional IPA threshold for confident activation (Z-score = 2.0), this result is interpreted as transcriptomic overlap with PE-related biology rather than evidence that Plac1 loss produces a PE-like disease state.

**Figure 8.**
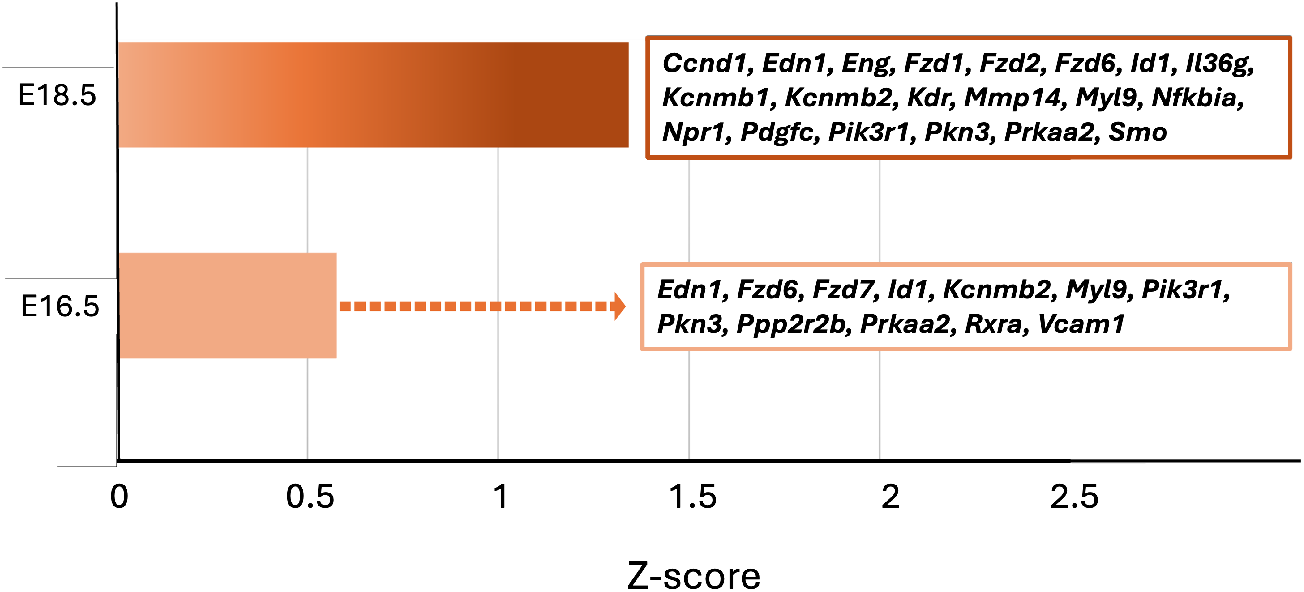
Transcriptomic Overlap with Preeclampsia-Related Pathway Features in Plac1-null Placentas. PE-associated pathways at each developmental age were identified using downregulated DEGs exhibiting at least a 1.5-fold decrease in expression compared to WT placentas. **E16.5** Solid light orange color. (Z-score = 0.58, FDR = 0.18) **E18.5** Dynamic orange fill depicting strengthening activation and significance level over time. (Z-score = 1.34, FDR = 0.02) **Text boxes:** DEGs populating the canonical PE pathway at each developmental age (Supplemental Tables S21 and S22). The vertical axis denotes gestational age. The horizontal axis denotes positive Z-scores. The dashed line (--->) represents the dynamic shift toward activation from E16.5 to E18.5

Unsupervised disease–gene enrichment analyses of E18.5 downregulated DEGs were performed using Enrichr disease-associated gene set libraries [78, 79] and provided additional context. DisGeNET identified vascular diseases, hypertensive diseases, dilated cardiomyopathy, vascular inflammation, and thrombosis among enriched disease categories (q = 0.01466–0.04304; thrombosis, q = 0.04180) [80]. By contrast, preeclampsia itself was identified only in the OMIM Expanded analysis but did not reach statistical significance (p = 0.0931, q = 0.1618; OR = 1.8) [81]. Together, these findings support transcriptomic convergence between the *Plac1*-null dataset and vascular, endothelial, inflammatory, and placental stress pathways relevant to PE, without defining a categorical PE transcriptional state.

Upregulated DEGs also included genes associated with fibronectin expression and glycosylation, molecular features relevant to PE biomarker biology. Elevated maternal serum glycosylated fibronectin (GlyFn) has been reported as a predictive marker for PE [82,83]. In the present dataset, placental *Fn1* expression was increased together with several genes involved in post-translational glycosylation, including members of the Galnt family, B4galnt2, St3gal3, St6gal1, and Mgat4a (**Table 6**). Because GlyFn assays rely on recognition of α2,6-sialylated epitopes, St6gal1 is relevant in this context. It should be noted, however, that altered GlyFn production, secretion, or maternal serum biomarker levels were not assessed in this study.

**Table 6.**
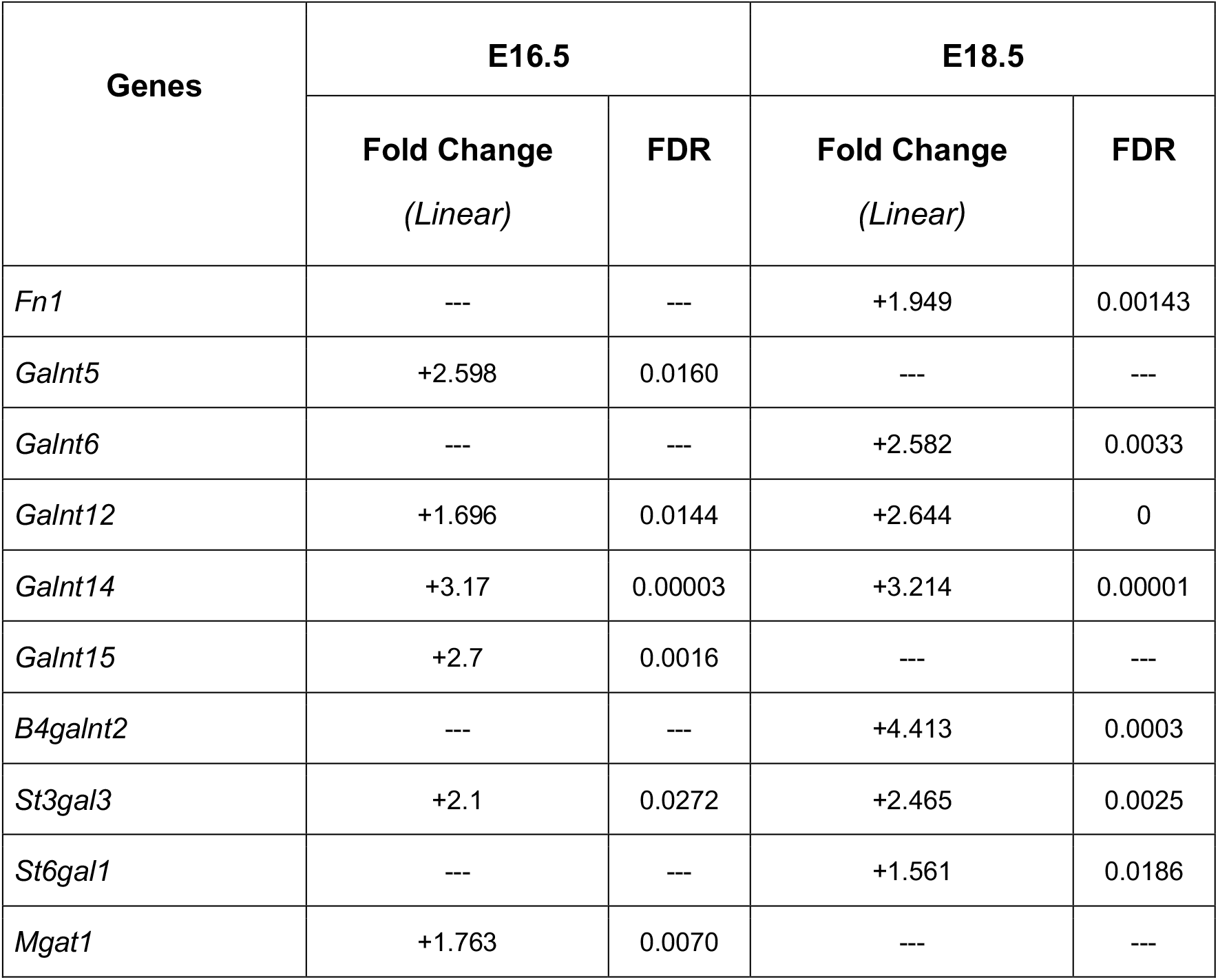
Upregulation of Fibronectin-associated Genes.

| Table 6. Upregulation of Fibronectin-associated Genes |  |  |  |  |
| --- | --- | --- | --- | --- |
| Genes | E16.5 |  | E18.5 |  |
|  | Fold Change<br>(Linear) | FDR | Fold Change<br>(Linear) | FDR |
| *Fn1* | --- | --- | +1.949 | 0.00143 |
| *Galnt5* | +2.598 | 0.0160 | --- | --- |
| *Galnt6* | --- | --- | +2.582 | 0.0033 |
| *Galnt12* | +1.696 | 0.0144 | +2.644 | 0 |
| *Galnt14* | +3.17 | 0.00003 | +3.214 | 0.00001 |
| *Galnt15* | +2.7 | 0.0016 | --- | --- |
| *B4galnt2* | --- | --- | +4.413 | 0.0003 |
| *St3gal3* | +2.1 | 0.0272 | +2.465 | 0.0025 |
| *St6gal1* | --- | --- | +1.561 | 0.0186 |
| *Mgat1* | +1.763 | 0.0070 | --- | --- |

#### 2.3.5 Predicted Upstream Regulator Patterns Are Concordant with Developmental and Stress-Associated DEG Signatures

To extend the pathway-level analyses, IPA “upstream regulator” analysis was used to identify predicted regulatory patterns associated with the downregulated and upregulated DEG datasets at E16.5 and E18.5 (Table 7). Because IPA upstream regulator analysis infers potential regulator activity from the expression patterns of curated downstream target genes, these results are interpreted as computational predictions rather than direct evidence of regulator activation or inhibition.

**Table 7.**
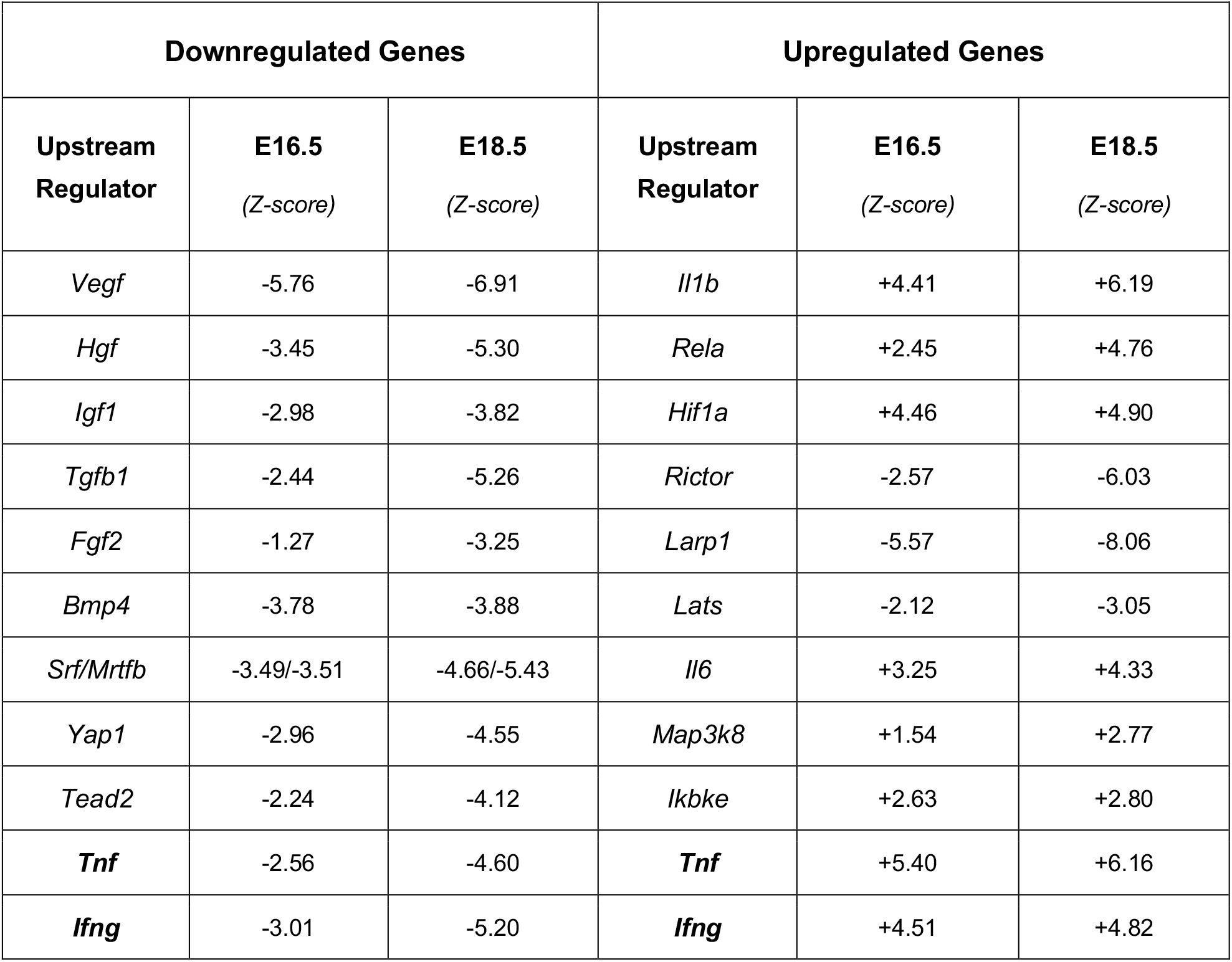
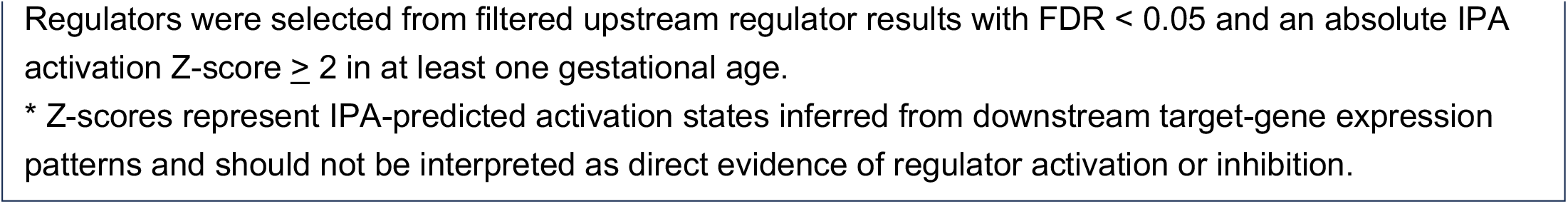
Representative upstream regulators illustrating opposing predicted activation states of vascular and developmental signaling versus inflammatory/stress-associated pathways in *Plac1*-null placentas.

| Downregulated Genes |  |  | Upregulated Genes |  |  |
| --- | --- | --- | --- | --- | --- |
| Upstream Regulator | E16.5<br>(Z-score) | E18.5<br>(Z-score) | Upstream Regulator | E16.5<br>(Z-score) | E18.5<br>(Z-score) |
| *Vegf* | -5.76 | -6.91 | *Il1b* | +4.41 | +6.19 |
| *Hgf* | -3.45 | -5.30 | *Rela* | +2.45 | +4.76 |
| *Igf1* | -2.98 | -3.82 | *Hif1a* | +4.46 | +4.90 |
| *Tgfb1* | -2.44 | -5.26 | *Rictor* | -2.57 | -6.03 |
| *Fgf2* | -1.27 | -3.25 | *Larp1* | -5.57 | -8.06 |
| *Bmp4* | -3.78 | -3.88 | *Lats* | -2.12 | -3.05 |
| *Srf/Mrtfb* | -3.49/-3.51 | -4.66/-5.43 | *Il6* | +3.25 | +4.33 |
| *Yap1* | -2.96 | -4.55 | *Map3k8* | +1.54 | +2.77 |
| *Tead2* | -2.24 | -4.12 | *Ikbke* | +2.63 | +2.80 |
| ***Tnf*** | -2.56 | -4.60 | ***Tnf*** | +5.40 | +6.16 |
| ***Ifng*** | -3.01 | -5.20 | ***Ifng*** | +4.51 | +4.82 |

Regulators were selected from filtered upstream regulator results with FDR < 0.05 and an absolute IPA activation Z-score $\geq 2$ in at least one gestational age.
\* Z-scores represent IPA-predicted activation states inferred from downstream target-gene expression patterns and should not be interpreted as direct evidence of regulator activation or inhibition.

Among downregulated DEGs, filtered upstream regulator results (absolute Z-score > 2; FDR < 0.05) included multiple predicted regulators associated with vascular development, growth-factor signaling, cytoskeletal organization, and developmental programs, including Vegf/Vegfa, Hgf, Igf1, Tgfb1, Fgf2, Bmp4, Srf/Mrtfb, Yap1, and Tead2 (Supplemental Tables S23 and S25). These predictions were broadly concordant with the GO, KEGG, and IPA canonical pathway findings described above, particularly the enrichment of membrane-associated signaling, Rho GTPase/cytoskeletal regulation, vascular adaptation, and developmental signaling pathways.

In contrast, upregulated DEG datasets were associated with predicted upstream regulators linked to inflammatory, cytokine, hypoxia-associated, and stress-responsive gene-expression patterns, including Il1b, Rela, Hif1a, Il6, Map3k8, Ikbke, Tnf, and Ifng (Supplemental Tables S24 and S26). These results were consistent with the enrichment of stress- and immune-associated pathways among upregulated genes. Several regulators, including Tnf and Ifng, appeared in both downregulated and upregulated DEG analyses with different predicted activation states, reflecting differences in the downstream target-gene subsets represented within each DEG group rather than uniform activation or inhibition of the regulator itself.

Together, the upstream regulator results provide an additional computational summary of the DEG patterns. Downregulated DEGs were associated with developmental, vascular, growth-factor, and cytoskeletal regulatory signatures, whereas upregulated DEGs were associated with cytokine, hypoxia-, and stress-associated signatures. These findings complement the GO, KEGG, and IPA canonical pathway analyses and are best viewed as hypothesis-generating computational summaries of the transcriptomic patterns. Complete upstream regulator results are provided in Supplementary Tables S23–S26.

### 2.4 Electron Microscopy (EM) of Plac1-null Placentas

TEM examination of placentas provided visual context for the molecular changes identified in the transcriptomic analyses. In the fields examined, structural differences were observed that were consistent with altered membrane organization, cytoskeletal regulation, and stress-response pathways. In an E18.5 WT placenta (**Figure 9A–C**), the labyrinth interhemal region displayed a well-organized trilaminar architecture. Sinusoidal trophoblast giant cells (sTGCs) lined the maternal blood space (MBS), and the syncytiotrophoblast (SynT) region formed an ordered layer overlying the basement membrane, followed by endothelial cells lining the fetal capillaries (FC). At higher magnification, mitochondria appeared small and relatively uniform, with discernible cristae. By contrast, in the *Plac1*-null placenta (**Figure 9D–F**), the corresponding interhemal region appeared less compact and less regularly organized in the field examined. The SynT region appeared less ordered, and enlarged mitochondria with less clearly defined cristae were observed in KO sTGCs.

**Figure 9.**
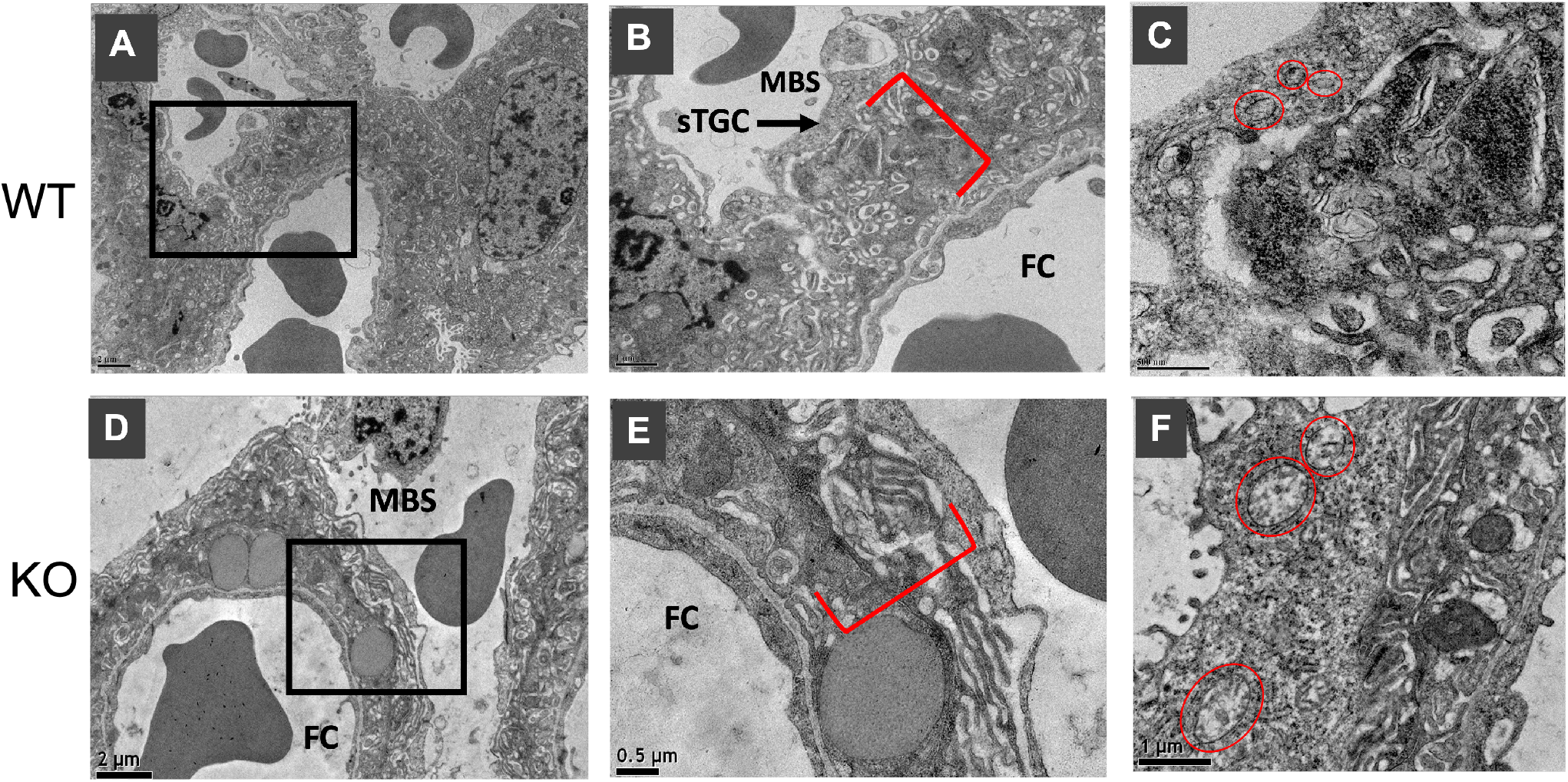
Transmission electron micrographs of the trilaminar interhemal region of E18.5 WT and KO placentas. WT Panels: The inset in panel **9A** shown at higher magnification in panel **9B**, depicts the trilaminar interhemal barrier separating the fetal capillaries (FC) from the maternal blood space (MBS). Sinusoidal trophoblast giant cells (sTGC; black arrow) are in direct contact with the MBS. A compact, well-defined region containing the syncytiotrophoblast (SynT) layers is indicated by the red brackets. Panel **9C** shows numerous small, well-defined mitochondria within an sTGC (red circles). KO Panels: Panels **9D** and **9E** show a similar interhemal region in the KO placenta. The SynT region appears more disordered and less compact. Panel **9F** depicts several enlarged mitochondria with poorly defined cristae (red circles). Scale bars = 0.5–2.0 µm.

These features are consistent with the broader transcriptomic pattern of altered membrane-associated signaling, cytoskeletal organization, and stress-response pathways in the *Plac1*-null placenta. However, because this analysis was limited to one placenta per genotype, these observations are presented as illustrative. Additional samples, systematic regional sampling, and quantitative ultrastructural analysis will be required to determine whether these features are reproducible and statistically associated with Plac1 loss. Source EM images are provided in Supplementary Figures S3–S8.

## 3. Discussion

This exploratory transcriptomic analysis of the *Plac1*-null placenta during late gestation identifies coordinated changes in gene expression associated with multiple biological processes important for placental organization and function. Integrating targeted gene-level curation of highly and reciprocally dysregulated genes with GO, KEGG, and IPA analyses, we identified recurrent themes involving membrane-associated signaling, cytoskeletal organization, vascular and maternal–fetal interface regulation, immune modulation, and stress-responsive pathways. These alterations occur in the context of the established *Plac1*-null placental phenotype and involve systems known to be important for normal placental development and pregnancy maintenance, providing a biologically coherent framework for understanding how the absence of *Plac1* may be associated with disruption of coordinated regulatory programs that evolve as the placenta matures. The internal consistency of the dataset is reflected in clear separation by genotype and gestational age, convergence across independent pathway analyses and concordance with the established *Plac1*null phenotype. Together, these features support the coherence of the major transcriptomic themes, while ultrastructural observations provide illustrative context. Because our transcriptomic window was restricted to E16.5–E18.5, alterations occurring earlier in placentation will require targeted investigation of those timeframes.

Membrane-associated signaling, actin-cytoskeletal regulation, and vascular-related processes were prominent features of the *Plac1*-null transcriptome. These systems are closely interconnected in the placenta, where trophoblast organization, maternal–fetal interface integrity, and labyrinthine vascular adaptation depend on coordinated signaling across membrane-proximal and cytoskeletal networks. The observed transcriptional changes in these categories are therefore consistent with disturbance of biological systems known to contribute to placental architecture and exchange function. Because the present study is based on transcriptomic associations rather than direct functional testing, these findings do not demonstrate that *Plac1* directly regulates any individual pathway or signaling node. Rather, they suggest that the loss of *Plac1* is associated with broad alteration of regulatory programs converging on processes essential for late-gestational placental maintenance.

Conservation of the *Plac1* sequence offers suggestive mechanistic context. *Plac1* shares ~30% homology with the zona pellucida binding protein ZP3 [1,84], a member of a conserved family of extracellular matrix proteins that organizes pericellular structures adjacent to the plasma membrane [84–86]. ZP3-like motifs are also present in membrane-associated extracellular glycoproteins including betaglycan, uromodulin, and glycoprotein 2 (GP-2) [87], and oocyte-enriched zona pellucida domain (ZPD) proteins (Oosp1–3) share homology with *Plac1* [88,89]. In *Drosophila*, ZPD proteins regulate epidermal cell shape and apical membrane–ECM interactions during embryogenesis [90], suggesting a conserved role in organizing membrane–cytoskeletal interfaces relevant to morphogenesis.

Specific *PLAC1*-protein interactions further support a membrane-adjacent regulatory role. PLAC1 interacts directly with desmoglein-2 (DSG2), a desmosomal protein [91], implicating roles in cell–cell adhesion, polarity, migration, and invasion. Dsg2 can also localize to apical compartments in enterocytes [92], raising the possibility of non-canonical roles linking the actin-rich terminal web to cytoskeletal systems. This is notable given human PLAC1 localization near the apical syncytiotrophoblast brush border and the F-actin-rich terminal web [5]. PLAC1 also binds extracellular FGF7/FGFRIIIb to activate AKT signaling [93,94] and interacts with the proprotein convertase furin to influence invasion-related Notch/NICD/PTEN signaling [95]. Together, these interactions converge on cytoskeletal regulation and Rho-linked signaling, consistent with the dominant pathway-level features observed in our transcriptomic analyses.

Although identified in cancer cell models, these interactions remain informative for developmental physiology. DSG2/Dsg2 is essential for cardiovascular development, and loss-of-function mutations are associated with cardiomyopathy and embryonic lethality in humans and mice [96–98]. The *Plac1*-null transcriptome displayed enrichment for Dsg2-associated cardiomyopathy programs, including dilated/arrhythmogenic cardiomyopathy, and both proteins are expressed in the developing myocardium. These observations raise the possibility that Plac1 contributes to Dsg2dependent cardiovascular function during critical developmental windows, although this remains a hypothesis requiring direct experimental testing. This possibility is also consistent with the observed 2.7-fold downregulation of Titin (*Ttn*) at E18.5 (Supplemental Table S6). Ttn provides structural scaffolding, passive elasticity, and mechanosensory function across the sarcomere [99], and truncating *Ttn* variants are the leading genetic risk factor for peripartum cardiomyopathy and have been identified in women with preeclampsia [100,101], implicating sarcomeric vulnerability as a shared feature of hypertensive pregnancy-associated cardiac disease. Because Dsg2 insufficiency produces Z-disk defects and impaired sarcomeric force transmission, disruption of the Plac1–Dsg2 axis during cardiomyocyte maturation may compromise desmosome–sarcomere coupling, providing a plausible conceptual framework for future studies of cardiomyopathic features.

As anecdotal context, the last surviving *Plac1* knockout male examined at 17 months exhibited marked cardiomegaly with histologic features of cardiomyopathic remodeling and secondary pulmonary congestion (Supplementary Figure S9). Because this observation was limited to a single animal, it is not used as evidence of a reproducible postnatal cardiac phenotype. Rather, it is retained as descriptive context supporting the rationale for future studies examining cardiac outcomes in larger cohorts with functional cardiac assessment.

*Plac1* interactions with membrane-associated proteolytic systems, including furin, represent a second mechanistic consideration. Furin is highly expressed in the syncytiotrophoblast, where it mediates proteolytic processing of substrates essential for placental differentiation and function [102,103]. Downregulation of the proprotein convertase *Bace2* at E18.5 in our dataset further suggests reduced capacity for regulated cleavage of membrane-associated substrates late in gestation (Supplemental Table S6). Because furin contributes to the maturation of multiple proteases within the secretory pathway, loss of *Plac1* may be associated with broader alterations in proteolytic processing through both transcriptional and post-translational mechanisms.

The previously reported increased risk of postnatal hydrocephalus in *Plac1* mutant mice provides developmental context for considering whether *Plac1*-associated membrane and proteolytic pathways have relevance beyond the placenta. Mutations in *L1CAM*, whose locus lies in close proximity to *PLAC1*, are the most common cause of X-linked hydrocephalus in humans, and hydrocephalus in *Plac1* mutant mice shows incomplete penetrance paralleling *L1cam*-mutant mouse models [104,105]. L1CAM function is regulated in part by proteolytic processing. Whether furin-dependent processing of substrates such as L1CAM operates similarly in neural tissue, where *Plac1* is also expressed, remains a plausible extension of these findings. However, the present placental transcriptomic data do not establish a neural mechanism.

Together, the interaction of Plac1 with both furin and Dsg2 places it at the intersection of membrane organization, cell–cell interaction, and regulated proteolysis in the syncytiotrophoblast. Because physiologically meaningful Plac1 interactions likely vary by cellular compartment, gestational timing, and differentiation state, defining these determinants across developmental contexts remains an important area for future investigation.

Among the canonical pathways identified by IPA, Rho GTPase signaling was among the most consistently downregulated, providing a plausible convergence point linking membrane-associated perturbations to cytoskeletal and structural placental programs. Rho-family GTPases coordinate extracellular signal transduction with actin organization, polarity, adhesion, migration, and vascular remodeling [106,107]. Additionally, signaling pathways that interact with Rho-associated machinery, including Wnt, TGFβ, VEGF, HGF, IGF, SHH, BMP, and PDGF, have established roles in placental and developmental biology [108–112]. Dysregulated tetraspanins observed in our dataset may further indicate altered tetraspanin-enriched microdomains, which organize integrins and growth factor receptors relevant to branching morphogenesis, angiogenesis, and ECM remodeling [113–116].

Dysregulated pathways were also enriched for annotations relevant to fetal organ systems where *Plac1* expression has been demonstrated, including cardiovascular, neurodevelopmental, kidney, lung, and musculoskeletal programs. These annotations are interpreted as shared developmental signaling programs rather than direct evidence of organ-specific pathology in the placenta or embryo. They are nevertheless consistent with epidemiological links between fetal growth restriction, prematurity, and developmental vulnerability [117,118], and with emerging concepts of placenta–brain and placenta–heart axes. Metabolic alterations, including disrupted selenoamino acid metabolism and thyroid hormone handling, offer additional plausible mechanisms by which placental dysfunction may influence fetal neurodevelopment.

Our findings also revealed transcriptional overlap with pathways and gene signatures associated with preeclampsia, most evident at E18.5, where pathway analyses identified increased representation of vascular, endothelial, inflammatory, and stress-responsive programs also implicated in preeclampsia-related placental dysfunction. Dysregulation of genes related to fibronectin processing and glycosylation is notable given the known association of glycosylated fibronectin to preeclampsia risk. Additionally, IPA disease-annotation analyses (Supplemental Tables S21, S22) identified cardiovascular and developmental categories overlapping with preeclampsia-related complications, particularly cardiovascular defects [119,120]. These findings indicate that the transcriptomic consequences of *Plac1* loss overlap with molecular features reported in preeclampsia and related placental stress states. We interpret this as consistent with broader convergence between the *Plac1*-null phenotype and pathways associated with placental dysfunction, while recognizing that these findings remain hypothesis-generating and require further validation in appropriately powered studies.

Preeclampsia remains a leading cause of perinatal mortality and long-term morbidity [121–123], with approximately half of susceptibility attributable to genetic factors across maternal, paternal, and fetal contributions [124,125]. Future studies should incorporate maternal phenotyping, earlier gestational windows, cell-resolved approaches, and inclusion of female heterozygous and knockout animals to define sex-specific responses. The progressive lethality observed across generations, including among heterozygotes [6], further suggests the possibility of intergenerational or epigenetic contributions. Consistent with this broader developmental framework, gene-level searches of the GWAS Atlas via Enrichr [78,79,126] identified significant *PLAC1* associations with birth weight, hematocrit, and waist circumference adjusted for body mass index (adjusted p-values = 0.0277 - 0.0352). Although these associations do not establish tissue-specific causality, they provide human genetic context linking *PLAC1* to traits broadly related to fetal growth, vascular and oxygen-transport biology, and later-life body-composition outcomes relevant to the DOHaD framework [127,128].

Taken together, these findings support the view that the *Plac1*-null placenta exhibits a transcriptional profile consistent with broad disturbance of coordinated placental support systems during late gestation, rather than isolated alteration of individual genes or pathways. The *Plac1*-null transcriptome is therefore most appropriately interpreted as a framework of molecular perturbations associated with an established placental phenotype rather than as evidence of direct mechanistic control by *Plac1* over any single affected pathway.

A central limitation of the present study is the limited biological replication inherent to this legacy dataset. The final DEG and pathway analyses were based on two biological replicates per genotype at each developmental stage. While the variance-modeling framework implemented in ExAtlas provides a structured approach for estimating gene-level variance across the dataset, it does not replace additional independent biological observations. Accordingly, the present findings should be interpreted as an exploratory, hypothesis-generating framework identifying coordinated expression patterns and pathway-level associations accompanying *Plac1* loss, rather than as definitive evidence of direct mechanistic regulation.

During preparation of this manuscript, Moreno-Irusta et al. [129] reported a contemporaneous, independent mechanistic analysis of PLAC1 function in rat and human trophoblast systems, including its interaction with furin and roles in trophoblast differentiation. These findings provide independent convergence with several principal features identified here, particularly the involvement of PLAC1 in membrane-associated signaling and trophoblast functional regulation. Conversely, the present study extends those observations by identifying systems-level transcriptional alterations across the intact placenta, placing PLAC1-associated biology within the broader context of maternal–fetal interface function and pregnancy-related disease pathways. Together, these complementary levels of biological organization provide a more integrated framework for understanding PLAC1 function across cellular, tissue, and developmental scales.

Finally, although this study focuses on the placenta, our findings parallel observations in cancer biology. *PLAC1* is reactivated in multiple malignancies where it associates with invasive/EMT-like transitions [130,131], proliferative and angiogenic signaling [93,94,132], and immunosuppressive microenvironments that mirror maternal–fetal immune tolerance [133,134]. In the context of minimal PLAC1 expression in normal adult tissues, these parallels support a model in which PLAC1 participates in conserved developmental programs that can be co-opted to similarly support cancer-related disease progression [10,135,136].

## 4. Conclusions

In summary, Plac1 loss is associated with coordinated transcriptomic changes in the late-gestation placenta, highlighted by altered placental developmental programs, reduced membrane-associated/Rho GTPase and actin-cytoskeletal signaling, and activation of immune, metabolic, and stress-response pathways. Together, gene-level curation and GO, KEGG, and IPA analyses are consistent with a model of developmental disequilibrium in the *Plac1*-null placenta, rather than disruption of a single linear pathway. These findings provide an exploratory, hypothesis-generating framework for future studies defining how Plac1 contributes to placental function, fetal developmental vulnerability, and pregnancy-related disease biology.

## 5. Materials and Methods

### 5.1 Mutant mouse model

The entire *Plac1* open reading frame (aa 2-173) was deleted in murine ES cells (C57BL/6NTac) as part of the NIH Knockout Mouse Program (KOMP). Blastocysts were injected with the *Plac1*-null ES cells and chimera obtained. After germline transmission was achieved the mice were bred against a C57BL/6 background (Jackson Laboratories) as previously described [6]. For the studies described in this report, timed matings between hemizygous, *Plac1* mutant (knockouts) or wild type (WT) males and heterozygous *Plac1* females (Hets) were carried out.

Pregnant females were sacrificed at E16.5 and E18.5 to obtain placental RNA in accordance with a protocol approved by the Institutional Animal Care and Use Committee (IACUC) of the University of South Florida-Morsani College of Medicine.

### 5.2 Genotyping and sex determination of mice

DNA was isolated from embryonic mouse tails using a DNeasy Blood and Tissue Kit (Qiagen). *Plac1* genotype was determined by PCR, using the primers:

5’-CCAATCATGTTCACCCACATTTCTAC-3 WT forward

5’-CCCTAAAAGAGCTATCATGGCATCT-3 Reverse

5’-GCAGCCTCTGTTCCACATACACTTCA-3 Neo universal forward

The cycling parameters were 94°C for 5 min followed by 10 cycles of 94°C for 15 s, 65°C for 30 s (decreased by 1°C at each repeat), and 72°C for 40 s; followed by 30 cycles of 94°C for 15 s, 55°C for 30 s, and 72°C for 40 s. PCR products were terminated with a final extension at 72°C for 5 min, then held at 4°C. A 1% agarose gel was used to visualize the generated wild type and mutant bands at 548 bp and 326 bp, respectively. Embryonic sex was determined by PCR using mouse SRY primers: 5’-TGGGACTGGTGACAATTGTC-3’ and 5’-GAGTACAGGTGTGCAGCTCT3’ [12] to score for maleness. The cycling parameters were 95°C for 4.5 min followed by 33 cycles of 95°C for 35 s, 55°C for 1 min and 72°C for 1 min, and final extension 72°C for 5 min, then held at 4°C. A 1% agarose gel was used to visualize the generated SRY fragment at 402 bp.

### 5.3 Microarray Analysis

Differential microarray analysis was carried out using the Agilent 4×44K mouse chip representing over 43,674 unique mouse transcripts as described by Carter, et al [137] and briefly summarized below.

#### 5.3.1 RNA Extraction, Target Labeling, Hybridization and Scanning

Total RNA was extracted from male WT and *Plac1*-null placentas at E16.5 and E18.5. The final microarray analysis included two biological replicates per genotype at each developmental stage: E16.5 WT males (n = 2), E16.5 *Plac1*-null males (n = 2), E18.5 WT males (n = 2), and E18.5 *Plac1*-null males (n = 2). RNA was extracted and purified using TriZol reagent (Invitrogen) per the manufacturer’s protocol. The quality and quantity of the preparations were assessed using an RNA 6000 Nano Lab-on-a-chip Kit with a 2100-Bioanalyzer system (Agilent Technologies).

Amplified cRNA labeled with Cyanine-3 CTP and Cyanine-5 CTP (Perkin-Elmer/NEN Life Sciences) was produced using a Fluorescent Linear Amplification Kit (Agilent Technologies) as specified by the manufacturer. The quality and size distribution of targets were determined by RNA 6000 Nano Lab-on-a-chip Assay (Agilent Technologies), and quantitated.

Fluorescent linear amplified cRNAs used in biological comparisons were then hybridized to Agilent 4×44K 60-mer oligo microarrays per the manufacturer’s instructions. Hybridized microarrays were washed according to the manufacturer’s protocol and scanned on an Agilent Technologies G2565AA Microarray Scanner System with SureScan technology.

#### 5.3.2 Data Processing and Statistical Analysis

Ratio data were extracted from scanned microarray images using Feature Extraction 5.1.1 software (Agilent Technologies). Dye-normalized, background-subtracted intensity and ratio data were exported to text and GEML-format files. Text output was originally processed using an application developed in-house (National Institute on Aging) to perform ANOVA analysis. Intensity values were filtered to remove values where probe error was greater than two times mean error and relative error was greater than 50%. Mean dye-swapped log(ratio) values were calculated, and mixed-model ANOVA was applied [137]. The potential for error variance was addressed by Bayesian adjustment to reduce false positives.

Subsequent to initial processing of the original scan data in 2012, the analytical platform was published online as the interactive tool ExAtlas [138] in 2015 and has been continuously updated as gene annotations evolve. The original scan data were reprocessed in 2026 using ExAtlas as previously described [138,139]. A comprehensive all-oligo matrix containing all probe features across genotypes was extracted from ExAtlas. To improve annotation reliability and reduce probe redundancy, the matrix was subjected to a two-step filtering process designed to retain the highest-quality and most reliable features. First, probes were filtered based on GenBank accession priority. For any given gene symbol, probes associated with curated RefSeq mRNA accession numbers (NM_) were preferentially retained, and all other accession types (XM_, AB_, AK_, etc.) were discarded if NM_accessions were present. In cases where no NM_accession was available, probes with predicted RefSeq accession numbers (XM_) were retained only if no alternative accession types were present; otherwise XM_probes were discarded in favor of other accessions (e.g., AB_, AK_). Second, among the remaining probes, a single best-performing oligo per gene was selected based on the highest F-statistic value (lowest p-value) from a one-way ANOVA across genotypes, ensuring retention of the probe with the greatest discriminatory power. This filtering process generated the best-oligo matrix (Supplemental Table S1), which served as the basis for construction of the ANOVA table (Supplemental Table S2) and all subsequent differential expression analyses between KO and WT placentas at E16.5 and E18.5. The female samples were excluded from the ANOVA because they lacked biological replication, thereby avoiding the contribution of non-replicated samples to the variance modeling applied to the final statistical analysis.

Statistical significance was derived using the variance-modeling framework implemented in ExAtlas, which estimates gene-level variance across the dataset rather than relying solely on gene-specific within-group replication. Because the final DEG analysis was based on two biological replicates per genotype at each developmental stage, the resulting gene lists are interpreted as exploratory and hypothesis-generating. Differentially expressed genes exhibiting at least a 1.5fold change and an FDR < 0.05 were subjected to KEGG, GO, and Ingenuity Pathway Analysis (IPA) to identify functionally relevant pathways. All data have been deposited in GEO (accession: GSE308499).

### 5.4 Transmission Electron Microscopy

Transmission electron microscopy was used to provide ultrastructural context related to Plac1 loss. To assess ultrastructural morphology, one E18.5 WT placenta and one E18.5 KO placenta were fixed overnight in 2.5% glutaraldehyde in 0.1 M phosphate buffer (pH 7.4) at 4°C. Samples were washed in 0.1 M sodium cacodylate buffer, pH 7.4. Post-fixation was performed in 1% osmium tetroxide in 0.1 M cacodylate buffer. After washing, the samples were dehydrated in a series of graded ethanol concentrations from 35% to 100%, followed by two washes in absolute acetone. The tissues were infiltrated and embedded in Embed812 resin. Ultrathin sections (80 nm) were mounted on copper grids and images were captured by a Gatan Orius digital camera mounted on a JEOL 1400 electron microscope at the Microscopy and Cell Imaging Core at the University of South Florida.

## Supplementary Materials

The following supplementary materials are available online for transparency and secondary analyses.

## Supplementary Tables

### Processed Expression Data and Statistical Analyses

- **Table S1.** Best-oligo-filtered gene expression matrix
- **Table S2.** Gene expression ANOVA results
- **Table S3.** Principal component analysis (PCA) output
- **Table S4.** E16.5 downregulated genes
- **Table S5.** E16.5 upregulated genes
- **Table S6.** E18.5 downregulated genes
- **Table S7.** E18.5 upregulated genes

### Gene Ontology (GO) Enrichment Analyses

- **Table S8.** GO analysis—E16.5 downregulated genes
- **Table S9.** GO analysis—E16.5 upregulated genes
- **Table S10.** GO analysis—E18.5 downregulated genes
- **Table S11.** GO analysis—E18.5 upregulated genes

### KEGG Pathway Enrichment Analyses

- **Table S12.** KEGG analysis—E16.5 downregulated genes (no significant Terms identified).
- **Table S13.** KEGG analysis—E16.5 upregulated genes
- **Table S14.** KEGG analysis—E18.5 downregulated genes
- **Table S15.** KEGG analysis—E18.5 upregulated genes

### Ingenuity Pathway Analysis (IPA)

- **Table S16.** IPA canonical pathways—E16.5 downregulated genes
- **Table S17.** IPA canonical pathways—E16.5 upregulated genes
- **Table S18.** IPA canonical pathways—E18.5 downregulated genes
- **Table S19.** IPA canonical pathways—E18.5 upregulated genes
- **Table S20.** IPA comparison analysis of shared canonical pathways (E16.5 and E18.5; Upregulated genes)
- **Table S21.** IPA summary—E16.5 downregulated genes
- **Table S22.** IPA summary—E18.5 downregulated genes
- **Table S23.** IPA Upstream Regulators – E16.5 downregulated genes
- **Table S24.** IPA Upstream Regulators – E16.5 upregulated genes
- **Table S25.** IPA Upstream Regulators – E18.5 downregulated genes
- **Table S26.** IPA Upstream Regulators – E18.5 upregulated genes

## Supplementary Figures

- **Figure S1.** Historical qRT-PCR plots for selected genes from the original microarray analysis
- **Figure S2.** IPA comparison analysis of shared canonical pathways (E16.5 vs. E18.5; upregulated genes)
- **Figures S3–S8.** Original electron micrographs corresponding to Figure 9A–F in the main manuscript
- **Figure S9.** Gross and microscopic images depicting cardiomegaly in an adult KO male

## Author Contributions

Conceptualization: Michael Fant, David Schlessinger and Ramaiah Nagaraja

Funding acquisition: Michael Fant and David Schlessinger

Resources: Michael Fant and David Schlessinger

Supervision: Ramaiah Nagaraja, David Schlessinger and Michael

Fant Investigation: Suzanne Jackman, Xiaoyuan Kong, Yulan Piao, and Michael

Fant Project administration: Ramaiah Nagaraja and Michael Fant

Methodology: Suzanne Jackman, Xiaoyuan Kong, Yulan Piao, Elin Lehrmann, Alexei Sharov, Ramaiah Nagaraja and Michael Fant

Data curation: Suzanne Jackman, Xiaoyuan Kong, Alexei Sharov, Elin Lehrmann, Andrew Varshine, Ramaiah Nagaraja and Michael Fant

Software: Elin Lehrmann, Andrew Varshine, Alexei Sharov, Michael Fant

Formal analysis: Alexei Sharov, Elin Lehrmann, Andrew Varshine, Ramaiah Nagaraja and Michael Fant

Validation: Xiaoyuan Kong

Writing – original draft: Michael Fant

Writing – review & editing: Alexei Sharov, Ramaiah Nagaraja, David Schlessinger, Suzanne Jackman, Xiaoyuan Kong, Yulan Piao, Andrew Varshine and Elin Lehrmann

## Funding

This work was supported in part by grants from the National Institutes of Health (HD-048862) and the March of Dimes (FY09503) awarded to MEF, as well as by the Intramural Research Program of the National Institutes of Health, which supported the microarray studies. The reanalysis phase (GO/KEGG enrichment and Ingenuity Pathway Analysis) was conducted using USF institutional core facilities. All associated core fees and IPA license access were paid from the corresponding author’s personal funds (MEF). Contributions by NIH-affiliated co-authors (DS, RN, YP, AS, AV, and EL) were performed as part of their official duties and constitute works of the United States Government. The content is solely the responsibility of the authors and does not necessarily represent the official views of the NIH or the U.S. Department of Health and Human Services.

## Institutional Review Board Statement

These studies required the use of a mutant mouse model. Approval was obtained from the IACUC at the University of South Florida who reviewed our breeding protocol, research strategy and plan to minimize any pain or discomfort in the animals. After careful review the project was approved on 06/18/2009 (IACUC# 3580R/3579M) and renewed on 05/02/2012 (IACUC# 4228R/4229M). Details of the IACUC practices and policies can be found at the following link: https://www.usf.edu/research-innovation/research-support/research-integrity-compliance/iacuc/

## Data Availability Statement

The raw and processed microarray data have been deposited in GEO and are openly available (accession: GSE308499). The curated microarray data summarized in this manuscript are included within the article and its Supplementary Materials. The original electron micrographs used in the manuscript are also provided in the Supplementary Materials. An earlier version of this work was posted on bioRxiv in April, 2026. DOI: https://doi.org/10.64898/2026.04.30.721637.

## Acknowledgements

The authors wish to thank Dr. Byeong Cha and Ms. Amanda Garces of the Microscopy Core Facility of the University of South Florida-Morsani College of Medicine (USF) for their assistance in performing the electron microscopy for these studies as well as Dr. Justin Gibbons of the USF Genomics Core for performing the GO and KEGG analyses. The authors also thank Professor Graham Burton (University of Cambridge, Cambridge, England) and Professor Robert Engelman (USF-Morsani COM/Moffitt Cancer Center) for reviewing the electron micrographs/tissue histology and providing valuable comments. The corresponding author (MEF) used Generative AI tools to brainstorm preliminary questions about gene and pathway functions and to provide editorial assistance regarding clarity and flow during manuscript writing. All information provided by these tools was verified against primary sources. All scientific interpretations and conclusions are those of the authors, and no Generative AI tools were used to determine data or figure content, generate figures, or conduct statistical analyses.

## Conflicts of Interest

The authors declare no conflicts of interest to report.

## Notes

### Competing Interest Statement

The authors have declared no competing interest.

### Summary of Updates

Figure 5 was replaced to correct a transcription error. Additional text was included to reference Table 6. References 7 and 9 were also renumbered to reflect their order of appearance in the manuscript. No changes were made to the Supplemental Material.

