## Supplementary material for "Transcriptomic Analysis of *Plac1* Ablation Reveals Broad Alterations in Signaling Pathways Essential for Prenatal Development and Overlap with a Preeclampsia-Associated Signature": 00_README_v4.1 copy.pdf

**README – Supplementary Materials**

This directory contains Supplementary Tables and Supplementary Figures associated with the manuscript *"Transcriptomic Analysis of Plac1 Ablation Reveals Broad Alterations in Signaling Pathways Essential for Prenatal Development and Overlap with a Preeclampsia-Associated Signature"*

**Supplementary Tables** include processed expression matrices **(Tables S1-S3)**, differential gene expression results **(Tables S4-S7)**, and GO **(Tables S8-S11)**, KEGG **(Tables S12-S15)**, and Ingenuity Pathway Analysis (IPA) outputs **(S16-S26)** referenced in the main text.

**Supplementary Figures** include historical RT–qPCR results **(Figure S1)**, IPA comparison visualizations **(Figure S2)**, original electron micrographs corresponding to Figure 9 in the manuscript (**Figures S3-S8)**, and images depicting cardiomegaly in an adult *Plac1*-null male **(Figure S9)**.

File names and indexing of all Supplementary Tables and Figures are provided below and in the Supplementary Materials section of the manuscript.

### File List:

Table_S1_Best_oligo_expression_matrix.txt

Table_S2_Anova_Differential_Expression_Plac1_KO_1-1.txt

Table_S3_PCA_output_Plac1_KO.txt

Table_S4_Genes_downreg_in_KOvsWT_malesE16.5_1.5fold_v1-1.txt

Table_S5_Genes_upreg_in_KOvsWT_malesE16.5_1.5fold_v1-1.txt

Table_S6_Genes_downreg_in_KOvsWT_malesE18.5_1.5fold_v1-1.txt

Table_S7_Genes_upreg_in_KOvsWT_malesE18.5_1.5fold_v1-1.txt

Table_S8_E16.5_GO_DownReg_Terms_Stats_Genes.tsv

Table_S9_E16.5_GO_UpReg_Terms_Stats_Genes.tsv

Table_S10_E18.5_GO_DownReg_Terms_Stats_Genes.tsv

Table_S11_E18.5_GO_UpReg_Terms_Stats_Genes.tsv

Table_S12_E16.5_KEGG_DownReg_Pathways_Stats_Genes.tsv ***(No Significant Terms)***

Table_S13_E16.5_KEGG_UpReg_Pathways_Stats_Genes.tsv

Table_S14_E18.5_KEGG_DownReg_Pathways_Stats_Genes.tsv

Table_S15_E18.5_KEGG_UpReg_Pathways_Stats_Genes.tsv

Table_S16_IPA_E16.5_DownReg_Canonical_Pathways_v1-1.txt

Table_S17_IPA_E16.5_UpReg_Canonical_Pathways_v1-1.txt

Table_S18_IPA_E18.5_DownReg_Canonical_Pathways_v1-1.txt

Table_S19_IPA_E18.5_UpReg_Canonical_Pathways_v1-1.txt

Table_S20_IPA_CompAnalysis_16.5x18.5_Upregulated_Canonical_Pathways_v1-1.txt

Table_S21_IPA_Summary_E16.5_DownReg_v1-1.txt

Table_S22_IPA_Summary_E18.5_DownReg_v1-1.txt

Table_S23_IPA_E16.5_Downregulated_Upstream Regulators_v1-1.txt

Table_S24_IPA_E16.5_Upregulated_Upstream Regulators_v1-1.txt

Table_S25_IPA_E18.5_Downregulated_Upstream Regulators_v1-1.txt

Table_S26_IPA_E18.5_Upregulated_Upstream Regulators_v1-1.txt

Figure_S1_Historical_qPCR_Results.pdf

Figure_S2_IPA_CompAnalysis_16.5x18.5_UpReg_Canonical_Pathways.pdf

Figure_S3_EM_Fig_9A_WT-PLB94-2b-4-6Kx.jpeg

Figure_S4_EM_Fig_9B_WT-PLB94-2b-5-15Kx.jpeg

Figure_S5_EM_Fig_9C_WT_50Kx-PLB94-2b-5-50Kx.jpeg

Figure_S6_EM_Fig_9D_KO_PLB94-7a maternal fetal interface-1 10k x.jpeg

Figure_S7_EM_Fig_9E_KO_PLB94-7a maternal fetal interface-1 30k x.jpeg

Figure_S8_EM_Fig_9F_KO_PLB94-7a maternal fetal interface-4-25k x.jpeg

Figure_S9_Cardiomegaly_Adult_KOxWT.pdf
