## Supplementary material for "Transcriptomic Analysis of *Plac1* Ablation Reveals Broad Alterations in Signaling Pathways Essential for Prenatal Development and Overlap with a Preeclampsia-Associated Signature": Figure_S1_Historical_qPCR_Results.pdf

**Figure S1.** Historical qRT-PCR plots for selected genes from the original microarray analysis.

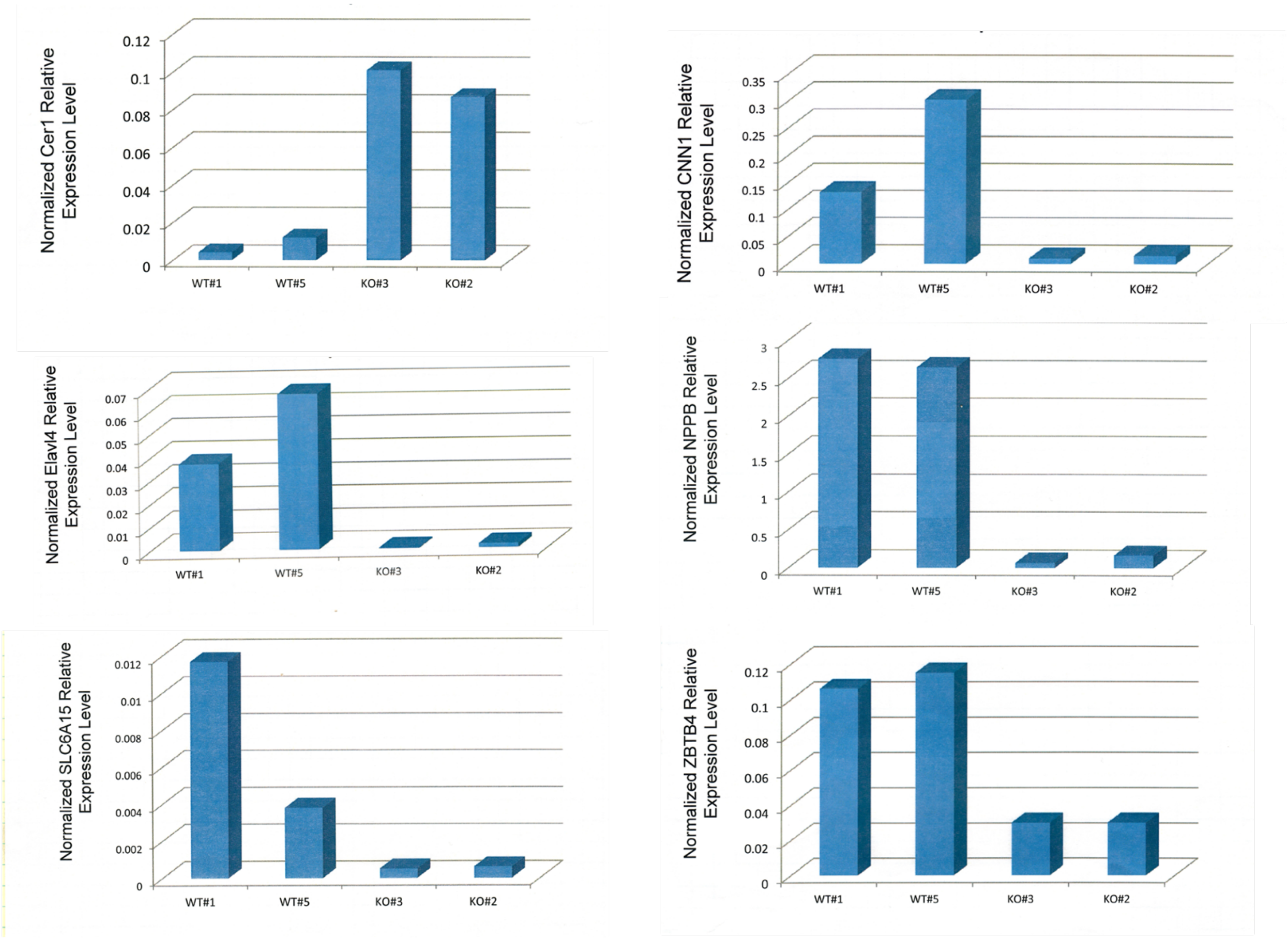


**Figure S1.**

Original-scale qRT-PCR plots for six genes assessed at E18.5 at the time of the original microarray analysis using TaqMan chemistry on cDNA generated from placental RNA, with 18S rRNA used as the reference control. These assays were performed using two biological replicates per genotype assayed in technical triplicate. Because the original raw qRT-PCR files and complete assay-performance documentation are no longer available, these data are provided only as qualitative historical observations of directional alignment with the microarray results. They are not used for statistical analysis or as independent validation of the transcriptomic dataset.
