## Supplementary material for "Transcriptomic Analysis of *Plac1* Ablation Reveals Broad Alterations in Signaling Pathways Essential for Prenatal Development and Overlap with a Preeclampsia-Associated Signature": Figure_S2_IPA_CompAnalysis_16.5x18.5_UpReg_Canonical_Pathways.pdf

**Figure S2.** Heatmap of IPA Canonical Pathways Associated with Upregulated Genes at E16.5 and E18.5

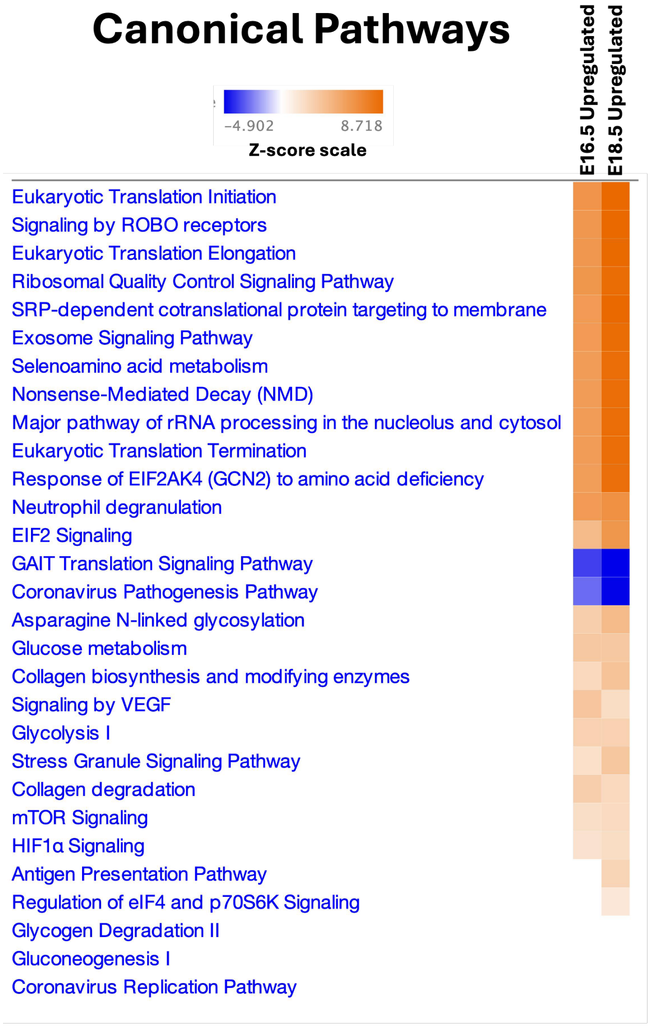


**Figure S2. Heatmap of IPA Canonical Pathways Associated with Upregulated Genes at E16.5 and E18.5**

Heatmap depicting the IPA “Comparison Analysis” of developmental stage-specific dysregulation of canonical pathways in KO placentas based on upregulated genes.

Color intensity reflects activation Z-scores: blue indicates negative Z-scores (predicted inhibition), and orange indicates positive Z-scores (predicted activation).

Pathways filtered for absolute Z-scores > 2 and Benjamini–Hochberg adjusted p-values < 0.05 at one or more developmental stages and ranked by absolute Z-score.
