## Supplementary material for "Transcriptomic Analysis of *Plac1* Ablation Reveals Broad Alterations in Signaling Pathways Essential for Prenatal Development and Overlap with a Preeclampsia-Associated Signature": Figure_S9_Cardiomegaly_Adult_KOxWT copy.pdf

### Supplementary Figure S9

#### Background

The last surviving male *Plac1* knockout (KO) mouse (17 months of age) died unexpectedly during off-site magnetic resonance imaging (MRI) performed to evaluate postnatal hydrocephalus, a known phenotype in *Plac1* mutant mice. An 11-month-old wild-type (WT) male was scanned concurrently as a control. Following death, necropsy was performed to obtain tissues for gross and histologic evaluation. Due to the unanticipated circumstances of death, tissue collection occurred under sub-optimal conditions.

During necropsy, a marked size discrepancy between the KO and WT hearts was immediately apparent. Gross photographs were obtained for side-by-side comparison under matched background and distance conditions (**Figure S9-1, Top Panel**). Additional images were acquired with each heart placed adjacent to its corresponding lungs prior to fixation (**Figure S9-1, Bottom Panel**). Tissue and body weights were not recorded.

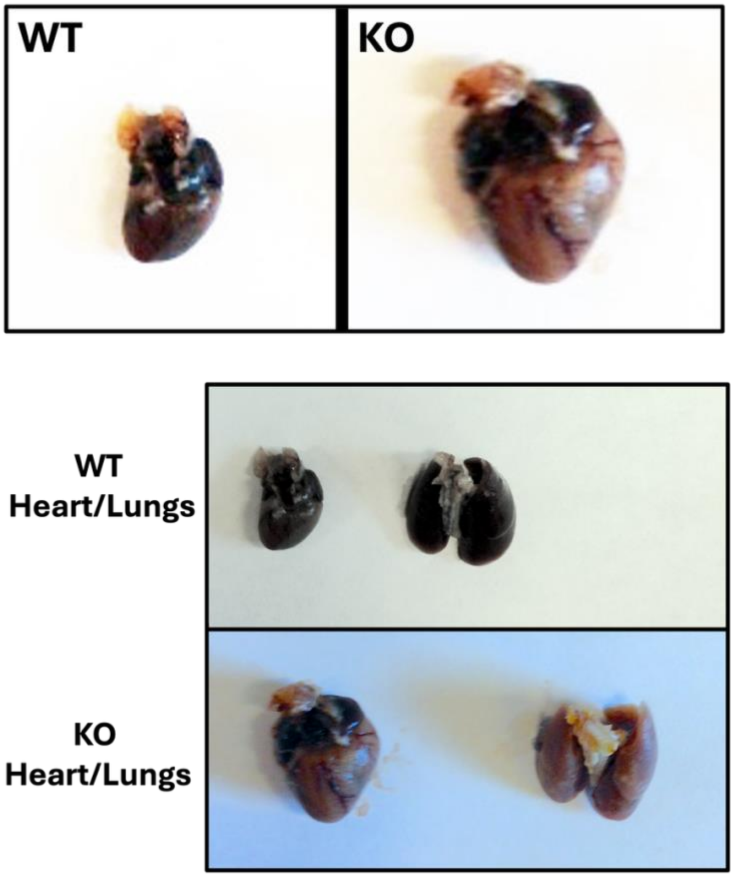


**Figure S9-1. Gross images of WT and *Plac1*-KO in adult mice.**
**Top Panel:** Side-by-side comparison of WT and *Plac1*-KO hearts demonstrating relative size differences.
**Bottom Panel:** Each heart positioned adjacent to its corresponding lungs prior to fixation.

**Histological Analysis**

Representative histologic sections of heart and lung tissue from both animals were prepared for histological evaluation (**Figure S9-2**) and independently reviewed by a board-certified veterinary pathologist at the University of South Florida Morsani College of Medicine and the Moffitt Cancer Center. The pathologist's findings are quoted below.

*"Heart of the Plac1 KO showed subgross evidence of marked cardiomegaly of undetermined cause. Hopefully staff obtained heart and body weights of mice so that organ/body weight ratios are known (although subgross images are convincing). If other Plac1 KO mice present with similar cardiomegaly, consider subgross dissection to help determine defect and cause (difficult to interpret from single section".*

*"Lung of the Plac1 KO mouse had vascular congestion and patchy consolidation (C) with eosinophillic fluid in alveolar spaces, moderate, mixed, often perivascular inflammatory infiltration (D), thickened edematous alveolar septa, and alveoli with numerous alveolar macrophages, all presumably referable to congestive cardiac insufficiency".*

*"Kidneys of the Plac 1 KO and WT mice lack significant abnormalities".*

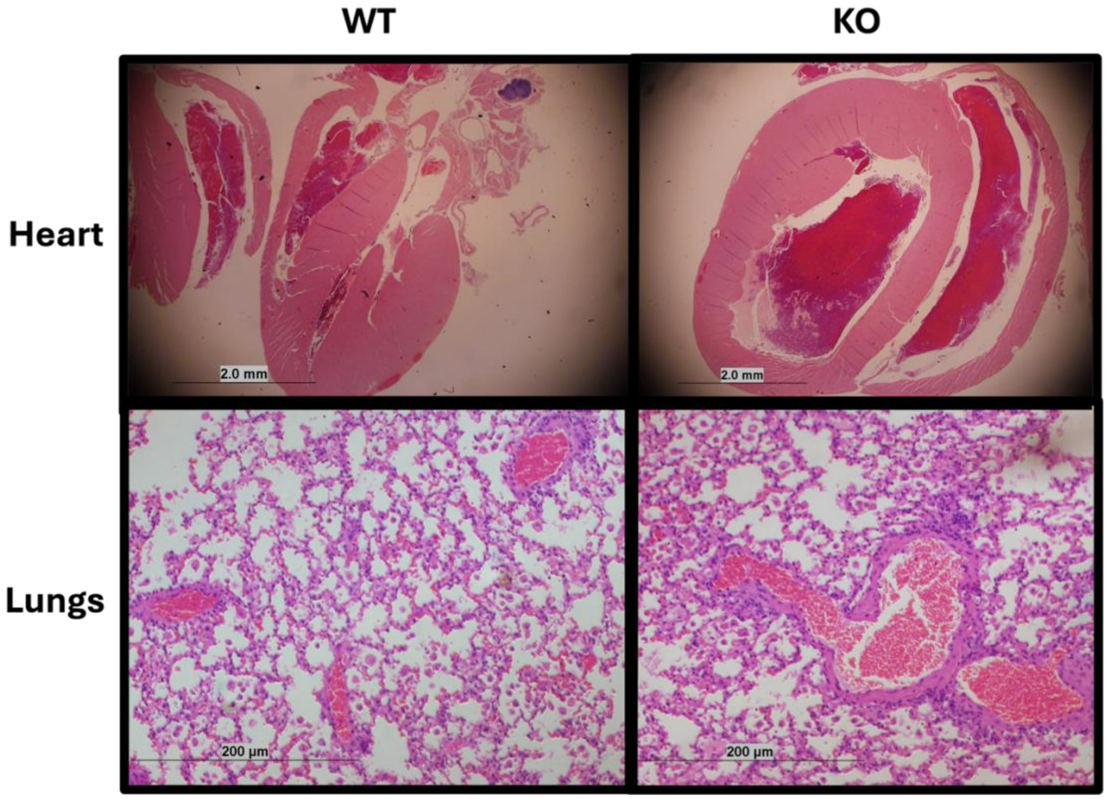


**Figure S9-2. Histological evaluation of WT and *Plac1*-KO heart and lung**

**Top Panels:** Sagittal heart sections from WT and *Plac1*-KO mice. Scale bar = 2.0 mm.
**Bottom Panels:** Lung tissue sections from WT and *Plac1*-KO mice. Scale bar = 200 µm.

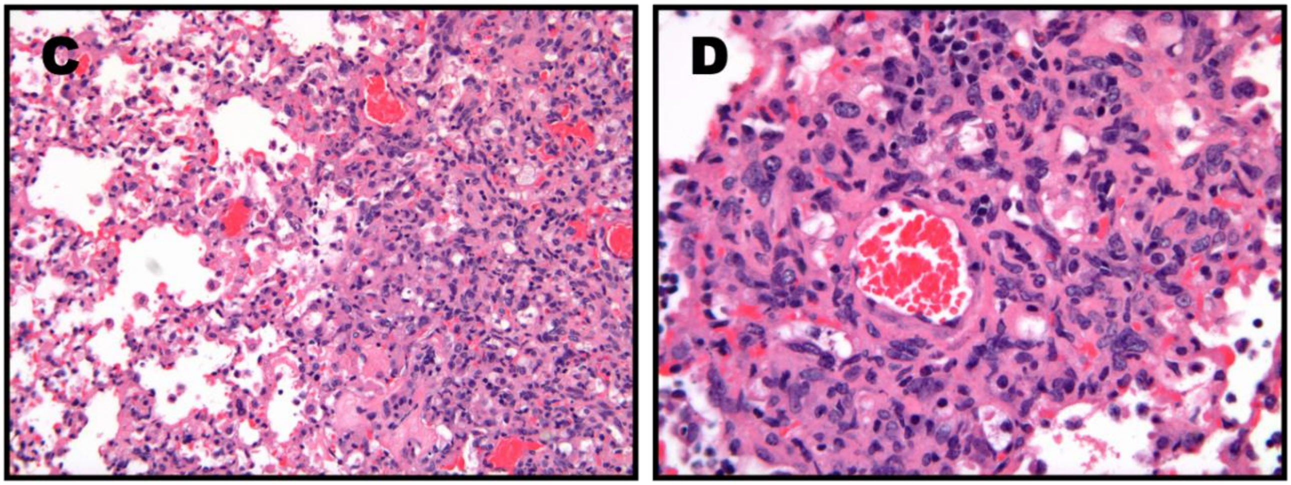


**Figure S9-3. Histological images of a *Plac1*-KO Lung** *(referenced by pathologist)*

**Panel C:** Image showing *patchy consolidation with eosinophillic fluid in alveolar spaces, perivascular inflammatory infiltration, thickened edematous alveolar septa, and alveoli with numerous alveolar macrophages,*
**Panel D:** Higher magnification of perivascular area.
