## Supplementary figures and images for "Transcriptomic Analysis of *Plac1* Ablation Reveals Broad Alterations in Signaling Pathways Essential for Prenatal Development and Overlap with a Preeclampsia-Associated Signature"

### Figure_S3_EM_Fig_9A_WT-PLB94-2b-4-6Kx copy.jpg

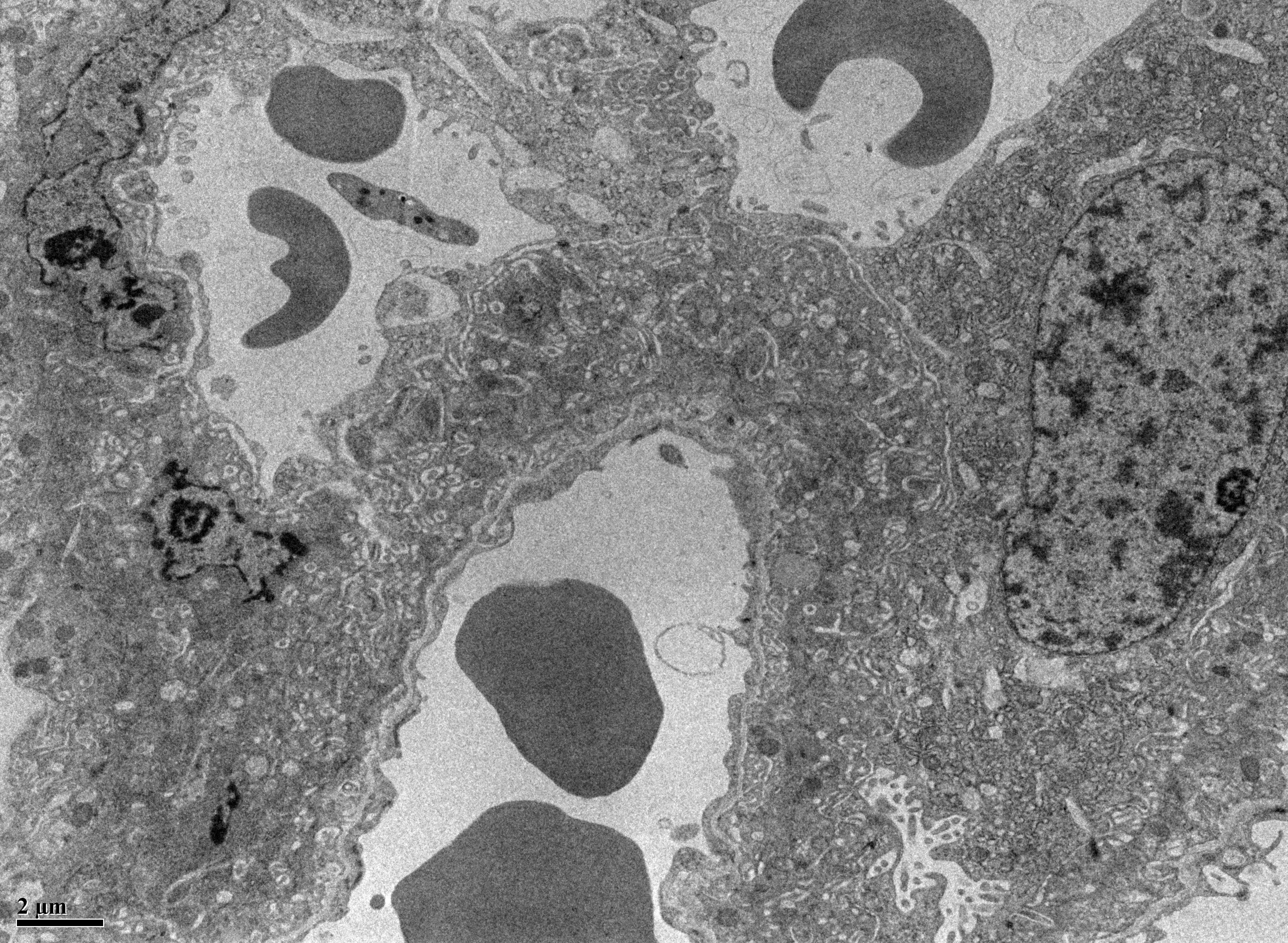

### Figure_S4_EM_Fig_9B_WT-PLB94-2b-5-15Kx copy.jpg

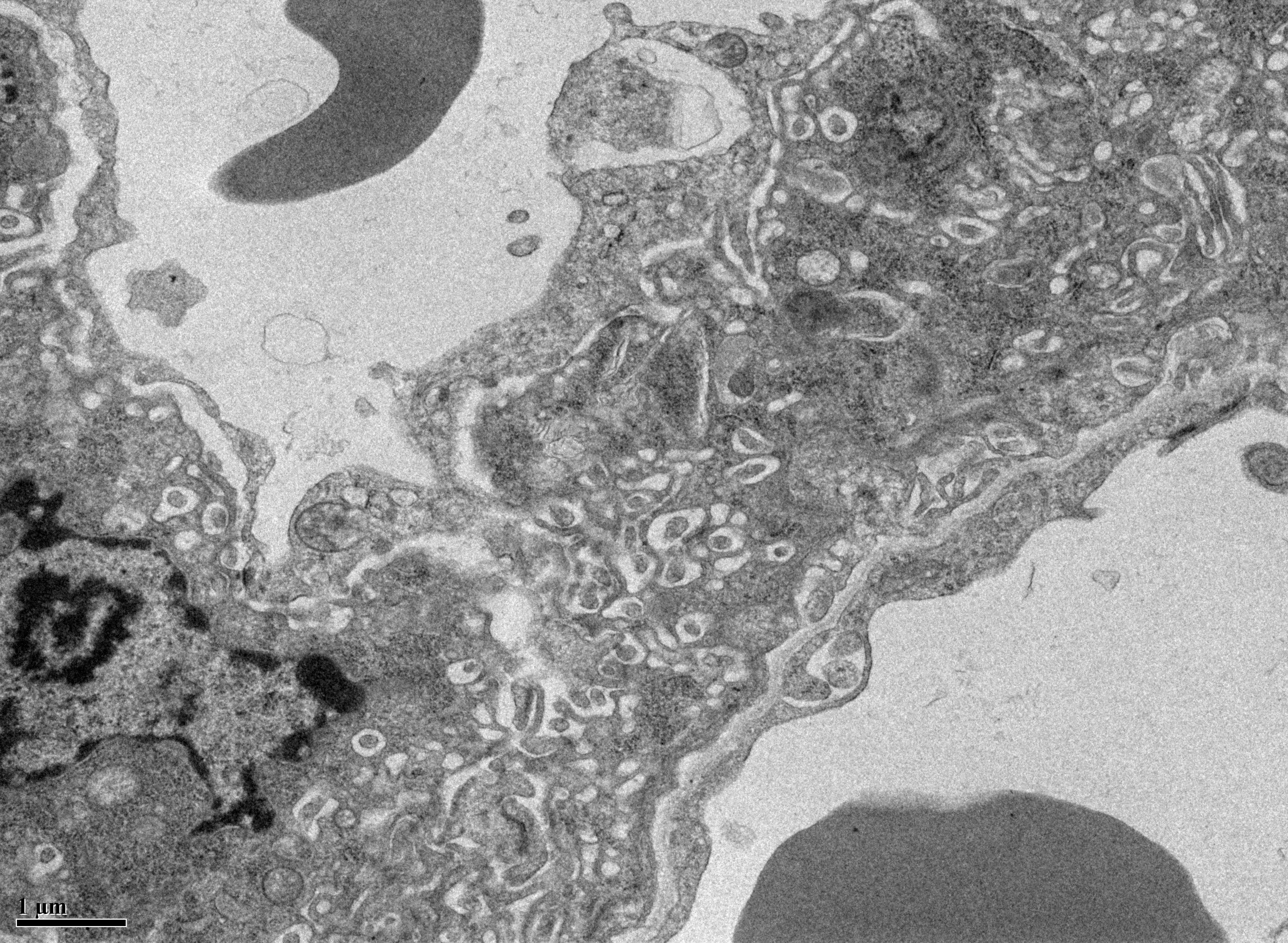

### Figure_S5_EM_Fig_9C_WT_50Kx-PLB94-2b-5-50Kx copy.jpg

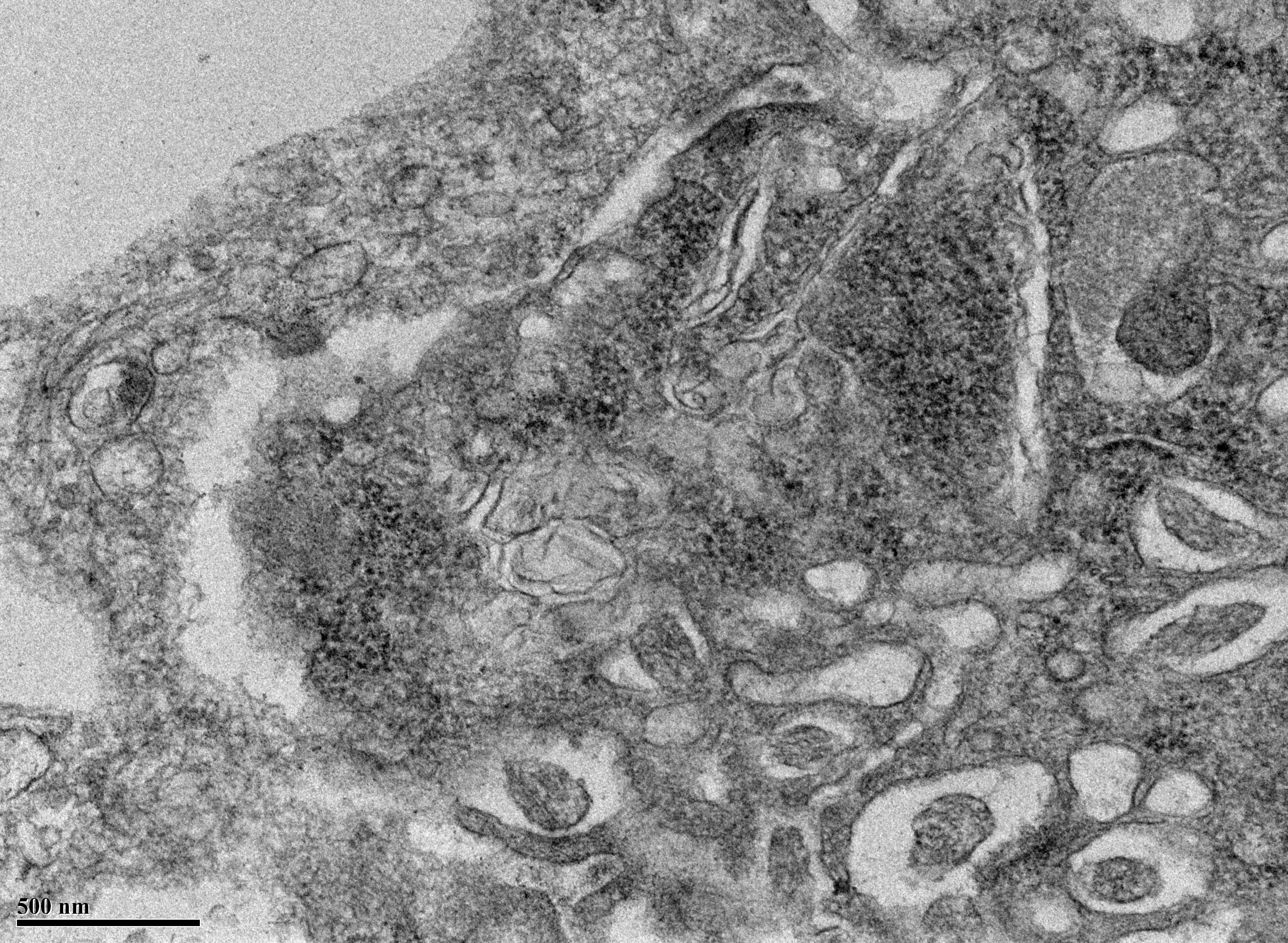

### Figure_S6_EM_Fig_9D_KO_PLB94-7a maternal fetal interface-1 10k x copy.jpg

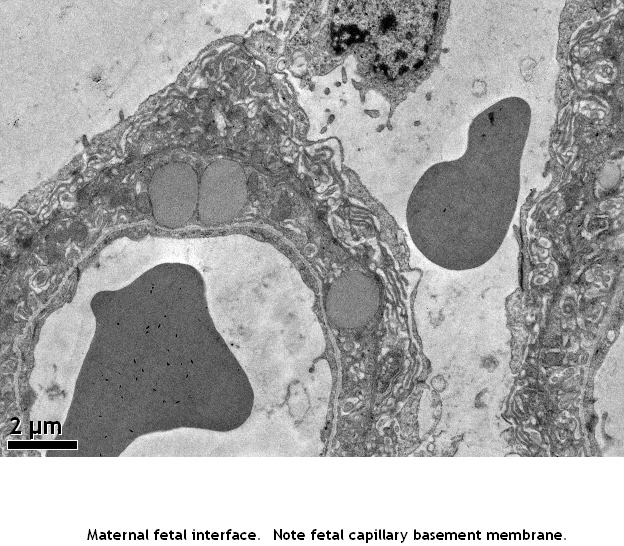

### Figure_S7-EM_Fig_9E_KO_PLB94-7a maternal fetal interface-1 30k x copy.jpg

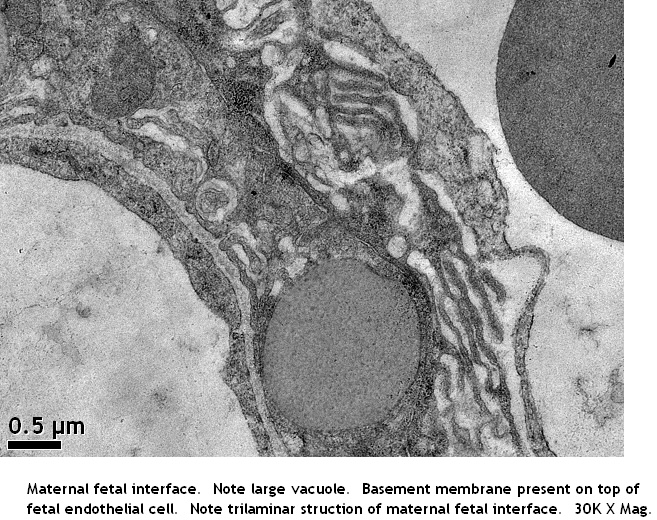

### Figure_S8_EM_Fig_9F_KO_PLB94-7a maternal fetal interface-4-25k x copy.jpg

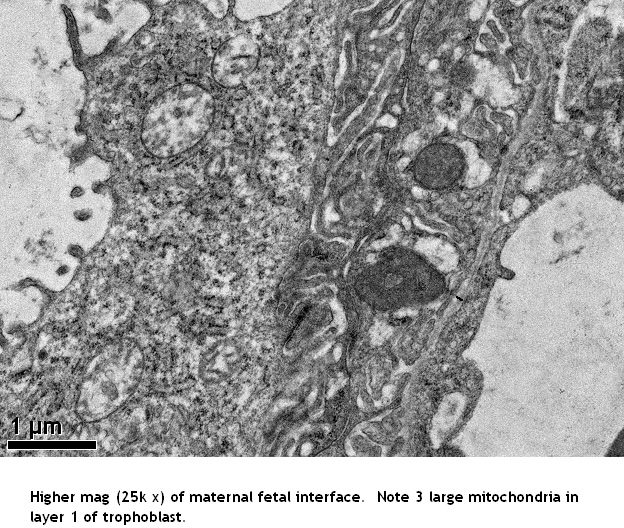
